# Glucose-dependent miR-125b is a negative regulator of β-cell function

**DOI:** 10.1101/2021.05.17.444559

**Authors:** Rebecca Cheung, Grazia Pizza, Pauline Chabosseau, Delphine Rolando, Alejandra Tomas, Thomas Burgoyne, Zhiyi Wu, Anna Salowka, Anusha Thapa, Annabel Macklin, Yufei Cao, Marie-Sophie Nguyen-Tu, Matthew T. Dickerson, David A. Jacobson, Piero Marchetti, James Shapiro, Lorenzo Piemonti, Eelco de Koning, Isabelle Leclerc, Karim Bouzakri, Kei Sakamoto, David M. Smith, Guy A. Rutter, Aida Martinez-Sanchez

**Author notes:** Authors contributed equally (Rebecca Cheung, Grazia Pizza). Corresponding author: Aida Martinez-Sanchez, PhD, Imperial College London, Du Cane Road, W12 0NN, London, UK.

## Abstract

Impaired pancreatic β-cell function and insulin secretion are hallmarks of type 2 diabetes. MicroRNAs are short non-coding RNAs that silence gene expression, vital for the development and function of β-cells. We have previously shown that β-cell specific deletion of the important energy sensor AMP-activated protein kinase (AMPK) results in increased miR-125b-5p levels. Nevertheless, the function of this miRNA in β-cells is unclear. We hypothesized that miR-125b-5p expression is regulated by glucose and that this miRNA mediates some of the deleterious effects of hyperglycaemia in β-cells. Here we show that islet miR-125b-5p expression is up-regulated by glucose in an AMPK-dependent manner and that short-term miR-125b-5p overexpression impairs glucose stimulated insulin secretion (GSIS) in the mouse insulinoma MIN6 cells and in human islets. An unbiased high-throughput screen in MIN6 cells identified multiple miR-125b-5p targets, including the transporter of lysosomal hydrolases *M6pr* and the mitochondrial fission regulator *Mtfp1*. Inactivation of miR-125b-5p in the human β-cell line EndoCβ-H1 shortened mitochondria and enhanced GSIS, whilst mice overexpressing miR-125b-5p selectively in β-cells (MIR125B-Tg) were hyperglycaemic and glucose intolerant. MIR125B-Tg β-cells contained enlarged lysosomal structures and showed reduced insulin content and secretion. Collectively, we identify miR-125b as a glucose-controlled regulator of organelle dynamics that modulates insulin secretion.

**Graphical abstract:** 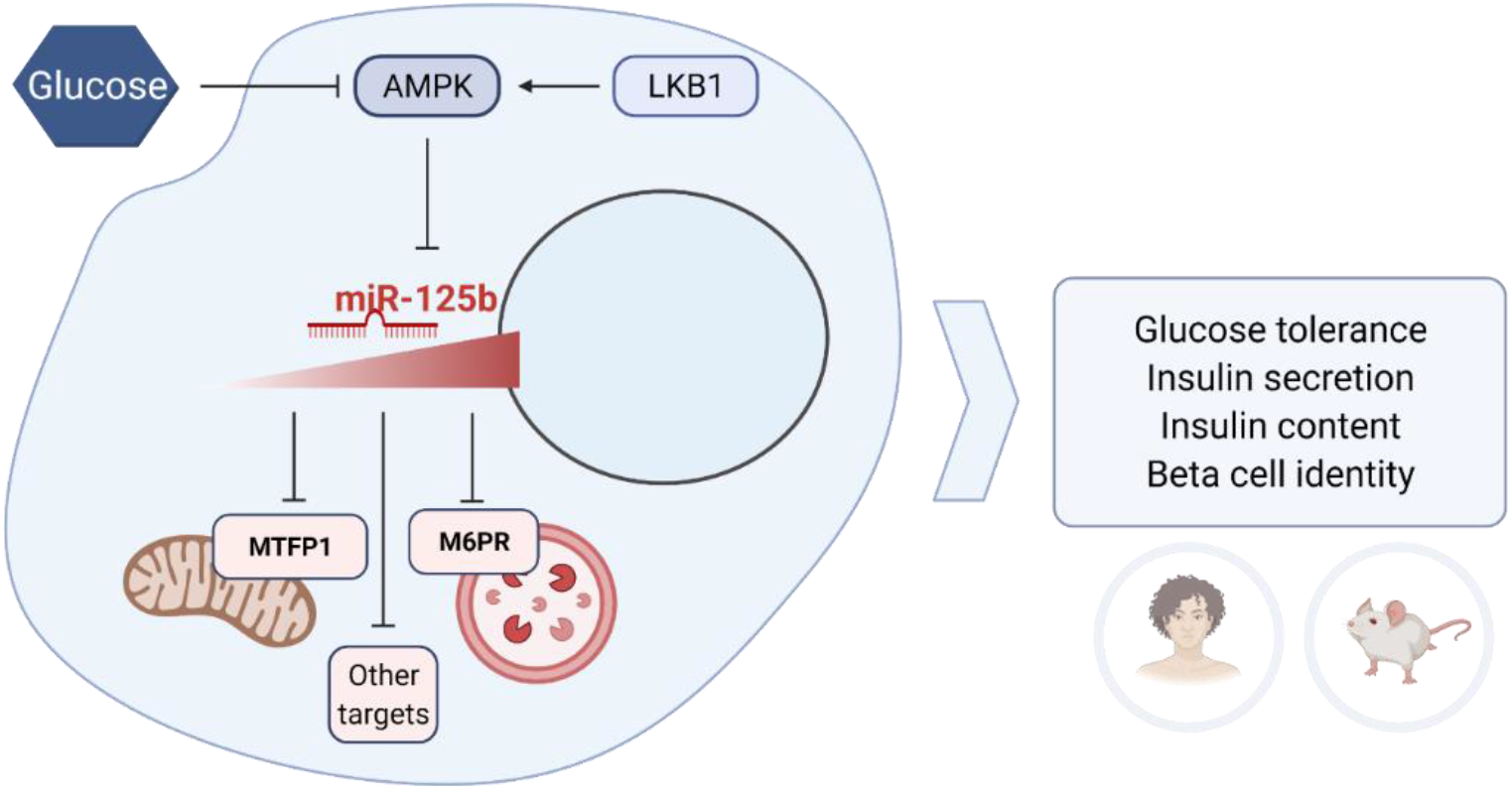

## INTRODUCTION

Pancreatic β-cells are essential regulators of glucose homeostasis, secreting insulin in response to increases in circulating levels of the sugar(1). Nevertheless, chronic hyperglycaemia has a negative effect on β-cell function and survival, contributing to the development of the type 2 diabetes(2).

MicroRNAs (miRNAs) are ~21 nucleotide non-coding RNAs that silence gene expression post-transcriptionally and are essential for endocrine cell development and function(3). Even though β-cells contain hundreds of different miRNAs, the function of only a few has been studied in detail.

MiR-125b (miR-125b-5p) is a highly conserved miRNA widely studied in the context of tumorigenesis as an important regulator of cellular differentiation and apoptosis(4). We have previously shown that miR-125b expression increases in islets after β-cell specific deletion of AMP-activated protein kinase (βAMPKdKO)(5), suggesting that AMPK may act as a negative regulator of miR-125b expression. AMPK activity is suppressed acutely in β-cells by high glucose and lowered in islets from subjects with T2D(6).

The above observations led us to hypothesize that inhibition of AMPK in response to high glucose would result in islet miR-125b up-regulation which may mediate some of the deleterious effects of hyperglycaemia in β-cells. Here we show that high glucose increases islet miR-125b expression and that miR-125b has an important negative effect in β-cell function both *in vitro* and *in vivo* by targeting genes involved in regulating lysosomal and mitochondrial function.

## RESEARCH DESIGN AND METHODS

### Cells and islets culture and transfection

MIN6 and EndoCβ-H1(7) cells and mouse and human islets were obtained, maintained and transfected with Lipofectamine 2000 (ThermoFisher Scientific) as described in Supplemental Methods.

### CRISPR-Cas9-mediated deletion of miR-125b in EndoCβ-H1 cells

EndoCβ-H1 cells were infected with lentiviral vectors expressing two gRNAs targeting *MIR125B-2* and a rat *Insulin* promoter (RIP)-driven hSpCas9. Lentivirus without gRNAs were used as control. Integrating EndoCβ-H1 cells were selected with puromycin. See Supplemental Methods for further details.

### Generation of transgenic mice

The cassette expressing miR-125b under the control of an rtTA-inducible promoter, was excised from pBI-LTet-MIR125B and used for pronuclear microinjection of C57BL6/J oocytes at the Centre for Transgenic Models, Basel (http://www.ctm-basel.ch/). A founder with a unique copy of the transgene was bred with mice expressing the rtTA under the control of the RIP7 promoter (RIP7-rtTA^+/+^). Mice bearing the transgene (MIR125B-Tg: RIP7-rtTA^+/−^, MIR125B Tg^+/−^) and controls (Control: RIP7-rtTA^+/−^) were used in subsequent experiments. Doxycycline (0.5 g/L) was continuously administered from the time of mating in the drinking water. Mice had free access to standard chow or, when indicated, ketogenic diet (Ssniff, Germany). *In vivo* procedures were approved by the UK Home Office Animal Scientific Procedures Act, 1986 (Licences PA03F7F0F/PP7151519.).

### RNA extraction, reverse transcription and qPCR

RT-qPCR was performed as previously described(5). For miRNA quantification, let-7d-3p (main figures) and miR-574-3p (supplemental figures) were used as endogenous controls due to their stable expression in our system(5). For mitochondria DNA/Nuclear DNA (mtDNA/nDNA) ratio analysis, islet DNA was extracted with 40ug/ml Proteinase K in SNET buffer.

### MiRISC immunoprecipitation, RNA sequencing library preparation, sequencing and analysis

MiRISC-immunoprecipitation was performed as previously described(8) with a mouse-anti-AGO2 antibody (clone E12-1C9, Abnova). Library preparation from mRNA enriched from total RNA with a NEBNext Poly(A) mRNA Magnetic Isolation Kit (NEB) and from miRISC-isolated RNA was performed using a NEBNext Ultra II Directional RNA Library Prep Kit from Illumina (NEB). Library preparation from human islets total RNA (200ng) was performed using a NEBNext Low Input RNA Library Prep kit (NEB).

Libraries were sequenced on a HiSeq4000, Mapping and differential expression analysis was performed with Salmon v1.3.0(9) and DESeq2(10).

For miRNA target identification, Total mRNA (T-RNA) and immunoprecipitated RNA (RIP-RNA) samples were treated as separate datasets and the two resulting gene lists of differential analysis were used to calculate the ratio of immunoprecipitated RNA to total mRNA (IP-RNA/T-RNA) for each gene.

Further details are provided in Supplemental Methods.

### Immunoblot and Immunohistochemistry

Western-immunoblot was performed with EndoCβ-H1 total protein (5-15μg) or 10-20 islets extract prepared with RIPA. Islets and slides from isolated pancreata for immunohistochemistry were prepared, visualized and quantified as previously described(11). Further details on the approaches used for blotting, β-cell mass, proliferation, apoptosis and cathepsin D/LAMP1 measurements, and an exhaustive list of all antibodies and kits used can be found in Supplemental methods.

### Electron microscopy

MIN6 and EndoCβ-H1 cells and whole isolated islets were fixed and prepared as in(12). Ultrathin 70-nm sections were examined in Tecnai Spirit or JEOL 1400 plus transmission electron microscopes. Images at 2000x magnification were analysed and quantified blindly in ImageJ as in(13).

### Intraperitoneal Glucose and Insulin tolerance tests (IPGTT, ITT) and *In vivo* Insulin secretion

Mice fasted overnight (IPGTT, Insulin secretion) or for 5h (ITT) were submitted to IPGTT or ITT following administration of 1g/kg body weight of glucose or 1.2U/kg insulin (NovoRapid, ITT), respectively, as previously described(14). For insulin secretion, mice were injected with 3g/kg glucose and blood insulin levels measured using an Insulin ELISA kit (Crystal Chem) following manufacturer’s instructions.

### Insulin secretion and content

Cell lines and islets were submitted to glucose or KCl stimulated insulin secretion as previously described(7; 14). Secreted and total insulin were quantified using an HTRF insulin kit (Cisbio) in a PHERAstar reader (BMG Labtech, UK) following the manufacturer’s guidelines. Insulin and protein content were measured by HTRF and BCA (Pierce), respectively, in RIPA cellular lysates or in full pancreas homogenates as previously described (11).

### Islet fluorescence imaging

Ca^2+^ and ATP imaging was performed in whole islets with Cal-520 (4.5μM; Stratech) and adenovirus-encoded Perceval probe, respectively, and TIRF was performed in dissociated islets infected with an adenovirus construct for NPY-venus as in(14; 15). Mitochondria morphology analysis was performed with Mitotracker Green™ (ThermoFisher) as in(15). See Supplemental Methods for further details.

### Whole-cell voltage-clamp electrophysiology

Voltage-clamp electrophysiology was performed in dissociated β-cells using patch electrodes and generating voltage-dependent Ca^2+^ (Ca_V_) channel currents through application of sequential 10 mV depolarizing steps as detailed in Supplemental methods.

### Extracellular flux analysis

Oxygen consumption rate was determined using the XFe96 Extracellular Flux Analyzer and a XFe96 FluxPak (Seahorse Bioscience) in groups of 3-6 size-matched islets/well embedded in Matrigel and incubated at 3 mmol/L glucose followed by 17 mmol/L glucose and oligomycin (5 μmol/L) following manufacturer’s instructions.

### Statistics

Statistical significance was evaluated with GraphPad Prism 9.0 software as indicated in the Figure legends. Correlation data was estimated using R software with ppcor package(16). All data are shown as means ± SEM. *p* < 0.05 was considered statistically significant unless otherwise specified.

### Study approval

All mouse in vivo procedures were conducted in accordance with the UK Home Office Animal (Scientific Procedures) Act of 1986 (Project licence PA03F7F0F to IL) and approved by the Imperial College Animal Welfare and Ethical Review Body. For human islet experiments, islets were isolated from the locations indicated in Sup Table 9 with full approval and informed consent.

### Data and Resource Availability

All data generated or analyzed during this study are included in the published article (and its online supplementary files). No applicable resources were generated or analyzed during the current study.

## RESULTS

### Glucose stimulates miR-125b expression via AMPK repression

To determine whether glucose regulates miR-125b expression in islets, we measured miR-125b in mouse and human islets cultured at increasing glucose concentrations. MiR-125b expression was significantly increased after culture at high (11-25mM) *vs* low (3.5-5.5 mM) glucose concentrations (Fig. 1A, S1A). Consistent with the possible *in vivo* relevance of these findings, expression of miR-125b in human islets was positively correlated with the body mass index (BMI) of donors, but not with their age or sex (Fig 1B, S1B,C).

**Figure 1.**
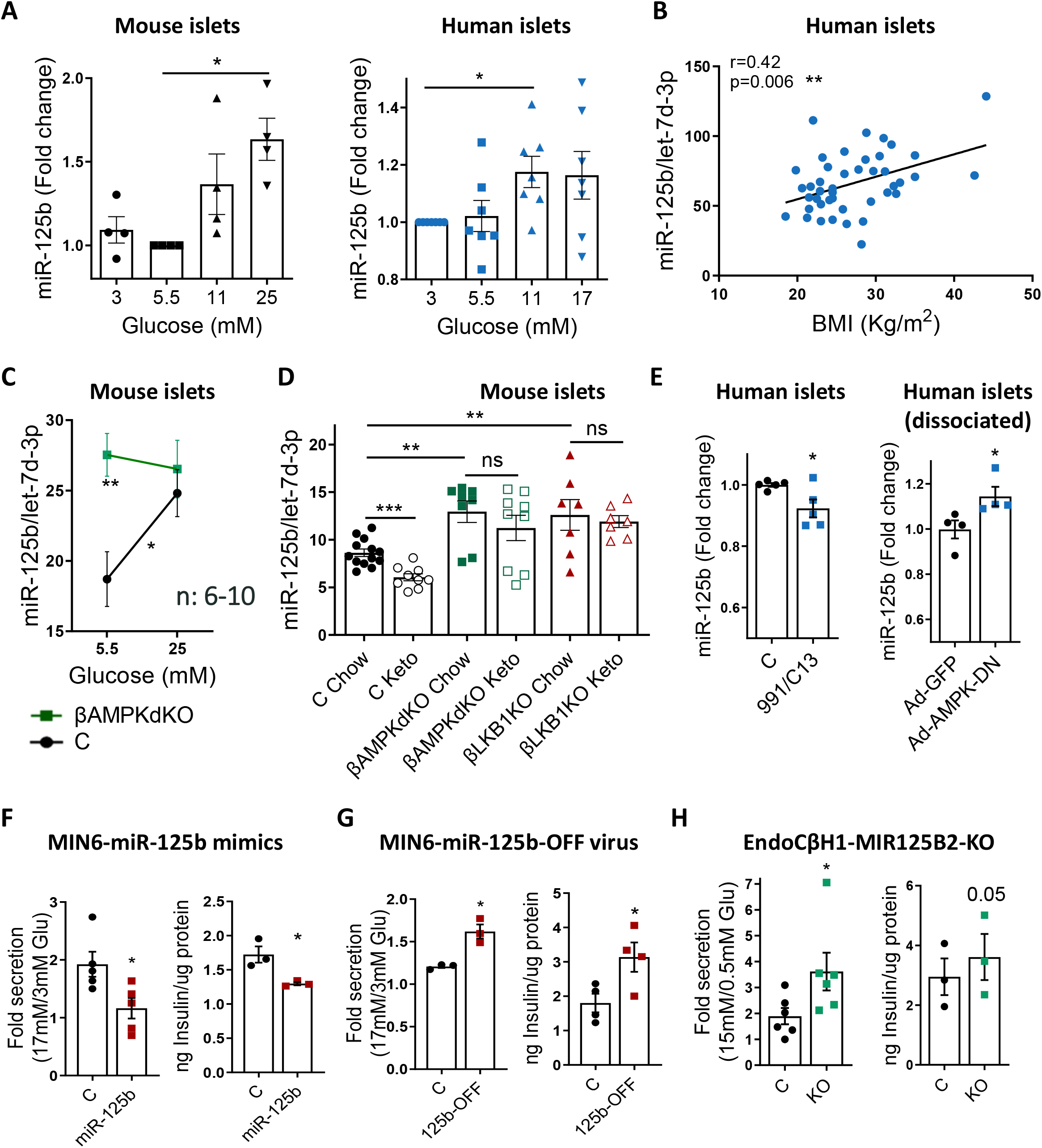
MiR-125b expression is stimulated by glucose via AMPK repression in islets and regulates insulin secretion in insulinoma/β-cell lines. RT-qPCR measurements of **A-E,** MiR-125b in (**A**) mouse and human islets cultured at the indicated glucose concentrations for 48h, (**B**) human islets (from non-diabetic donors) kept for 24h at 5.5mM glucose plotted against donor’s BMI, (**C**) islets from βAMPKdKO (green) and control (C, black) mice cultured at 5.5 or 25mM Glucose for 48h, (**D**) islets from βAMPKdKO (green), βLKB1KO (red) and control (C, black) male and female mice fed a chow or a ketogenic (Keto) diet for 28 days and (**E**) human islets treated with the AMPK activators C-13 (50nM) and C-991 (20nM) for 16h (left hand-side) and dissociated human islets infected with adenovirus expressing a dominant-negative AMPK protein (Ad-AMPK-DN) or empty control (Ad-GFP) at 5 MOI for 48h (right hand-side panel). Each dot represents islets from a single mouse or human donor. MiR-125b expression is normalized to that of the endogenous control let-7d-3p (4). **F-H** Glucose-stimulated insulin secretion (GSIS, left hand-side panels) and insulin content (righ hand-side panels). GSIS was quantified after 30 minutes of 17 or 15 mM glucose stimulation following overnight preincubation at 3 mM glucose of MIN6 (**F, G)** and EndoCβ-H1 with CRISPR-mediated miR-125b knockout (**H).** MIN6 cells were transfected with 5nM miR-125b (125b) or control (C) mimics (**F**) or infected with adenovirus expressing a miR-125b inhibitor (125b-OFF) or a non-targeting control (C) at 10 MOI (**G**) 48h before the experiments. Intracellular protein content was quantified by BCA to calculate insulin content per μg of total protein. Data is presented as fold change of basal level. Each dot represents an independent experiment with two-three technical replicates. Experiments were performed with three different populations of CRISPR-mediated miR-125b KO and control cells. Error bars represent SEM. ns=no significant, \**p* <0.05, \*\**p* <0.01, \*\*\**p* <0.001, one-way ANOVA (repeated measures) and Dunnet multiple comparisons test (A), Pearson Correlation adjusted by age and sex (B), two-way ANOVA (repeated measures) and Bonferroni multiple comparisons test (C), unpaired Welch (D) and paired Student (E-H) *t* test.

Islets from βAMPKdKO mice displayed significantly higher miR-125b levels than controls when cultured at a low (5.5mM) glucose concentration (Fig. 1C, S1D) whereas miR-125b expression remained unchanged in islets cultured at high glucose (25mM). To determine whether this glucose/AMPK-dependent regulation of miR-125b also occurred *in vivo*, we fed control, βAMPKdKO or βLKB1KO (mice with a β-cell specific deletion of the AMPK-upstream kinase Liver kinase B1, LKB1/STK11(17)) animals a ketogenic (low sugar) diet. As previously showed(5), βAMPKdKO islets displayed higher levels of miR-125b than controls and the same was observed in βLKB1KO islets (Fig. 1D). Feeding a low-sugar diet resulted in a significant decrease in islet miR-125b in control animals but not in islets from βAMPKdKO and βLKB1KO mice (Fig. 1D). Moreover, culture of human islets with the selective AMPK activators C13 and 991(5) or with adenovirus encoding a dominant-negative form of the enzyme caused a small, but significant, reduction and increase, respectively, in miR-125b expression (Fig. 1E, S1E). Taken together, these results indicate that high glucose stimulates miR-125b expression in both mouse and human islets by inhibiting AMPK activity.

### MiR-125b regulates insulin secretion in β-cell lines

The most important characteristic of the β-cell is its capacity to respond to high levels of circulating glucose by secreting insulin(1). Glucose-stimulated insulin secretion (GSIS) in MIN6 cells with ~40-fold increased miR-125b (Fig. S2A) following transient transfection with miR-125b mimics was lower than in control cells and these cells contained 25% less insulin (Fig. 1F, S2B). Conversely, inhibition of miR-125b function with an adenovirus encoding for a miR-125b inhibitor resulted in increased GSIS and insulin content (Fig 1G, S2C).

To explore whether miR-125b regulated insulin secretion in a human β-cell line we next generated human EndoCβ-H1 cells(18) with miR-125b loss-of-function using CRISPR-Cas9 (EndoCβH1-MIR125B2-KO, Fig S3A). EndoCβH1-MIR125B2-KO cells contained ~80% less mature miR-125b than controls (Fig. S3B), showed similar viability (Fig. S3D) and, as anticipated, secreted more insulin in response to high glucose than controls, although only a small, non-significant, increase was detected in insulin content (Fig. 1H, S2D).

### A high-through approach identifies miR-125b target genes

To further explore the genes and molecular pathways regulated by miR-125b, as well as its mechanism of action, we sought to explore the impact of the miRNA on β-cell gene expression and to identify direct targets in a high-throughput manner. MiRNAs guide the miRNA-induced silencing complex (miRISC), including an Argonaute protein to the target mRNAs which results in mRNA destabilization and/or inhibition of translation(19). Thus we used an experimental approach that combined overexpression of miR-125b in MIN6 cells, immunoprecipitation of miRISC/target mRNAs using an antibody against AGO2 and high-throughput sequencing and differential analysis of cellular total and immunoprecipitated RNAs(20) (Fig. S4A). As a consequence of miR-125b overexpression, we expected miR-125b targets to be reduced or unchanged (if repression occurs exclusively at the level of translation) in the total mRNA (T-RNA, Table S1) fraction and increased in the precipitated (RIP-RNA, Table S1) fraction. Thus, for miR-125b direct targets we expect a RIP-RNA to T-RNA ratio > 1 following miR-125b overexpression. To validate our approach we sorted all genes according to RIP-RNA/T-RNA ratio (Table S1) and performed unbiased motifs enrichment analysis using cWords(21). As anticipated, cWords found a significant enrichment of miR-125b seed-matching sequences in both the 3’ non-translated (3’UTR) and coding (CDS) sequences of highly ranked (high RIP-RNA/T-RNA ratio) genes (Fig. S4B), suggesting that miR-125b-mRNA target interactions occur through both these regions. 180 mRNAs were enriched in miRISC following miR-125b overexpression with an RIP-RNA/T-RNA > 1.5 (Fig. 2A, Table 1). Out of these, 60 were significantly down-regulated at the mRNA level following miR-125b overexpression (Table 1, padj < 0.1) and 87 contained predicted binding sites in their 3’UTRs according to TargetScan, ~50% (49) of which are conserved in humans (Table 1).

**Figure 2.**
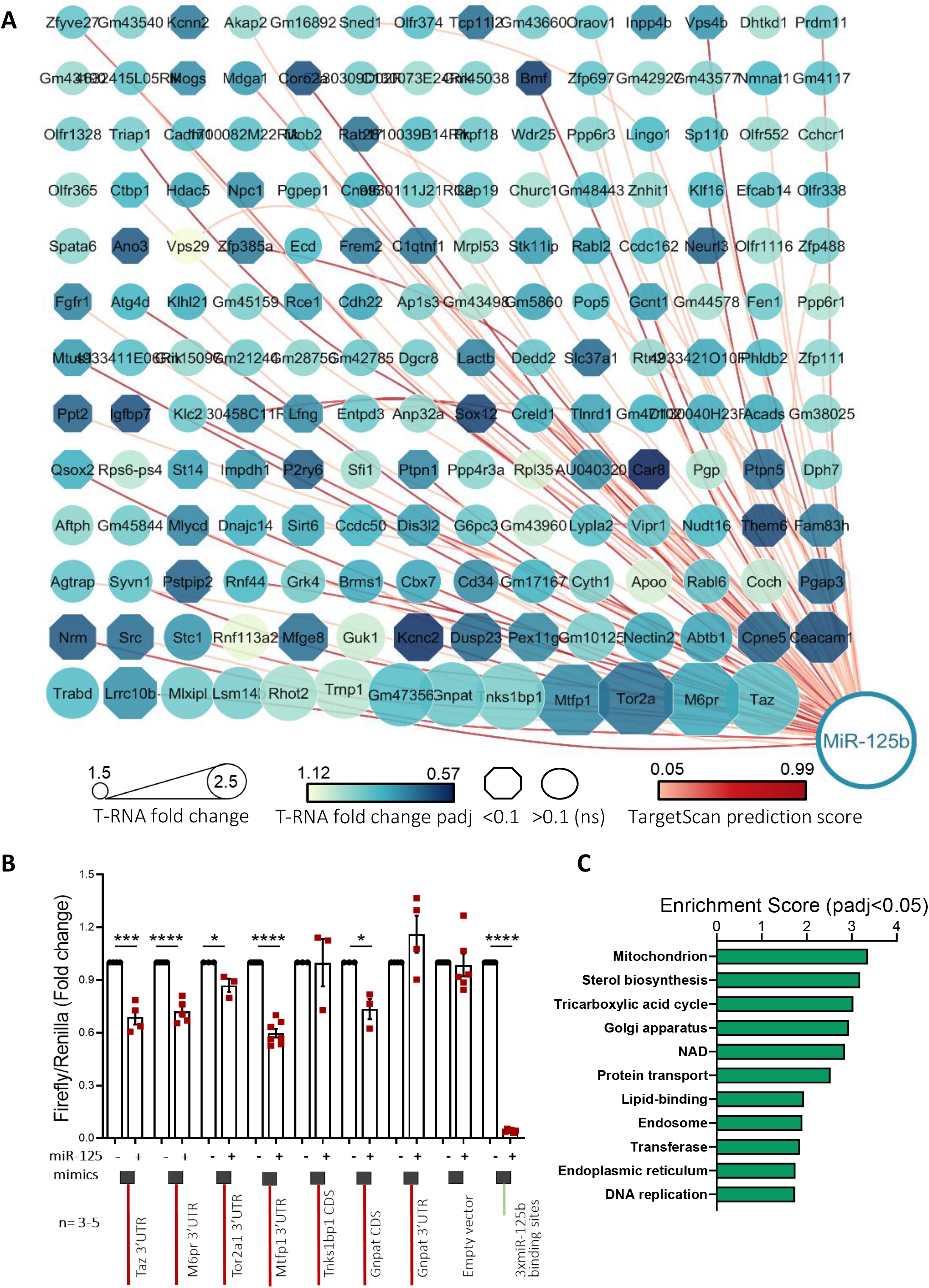
High-through identification and validation of miR-125b target genes. MIN6 cells were transfected with 5nM control or miR-125b mimics **A)** Cytoscape(49)-generated layout of genes with RIP-RNA/T-RNA ratio > 1.5 upon miR-125b overexpression (arbitrary cut-off) in MIN6 cells were transfected with 5nM control or miR-125b mimics. Node size represents the extent of the RIP-RNA/T-RNA ratio (the larger, the higher). The intensity of the gene node colour indicates the fold downregulation of each gene upon miR-125b expression (T-RNA). Downregulation at the transcript level was statistically significant for those genes represented in an octagonal shape. Lines connecting the nodes/genes with miR-125b indicate the presence of a predicted binding site for miR-125b as identified by TargetScan, with the intensity in the colour of the line representing the score (Total context score). **B)** Validation of miR-125b binding to the regions (full-length CDS or 3’UTR, as indicated) containing predicted miR-125b binding sites of six identified targets with highest RIP-RNA/T-RNA. Luciferase reporter assay of MIN6 cells cotransfected with pmirGLO plasmids containing the indicated target downstream the luciferase ORF and miR-125b (+) or non-targeting (-) control mimics. Samples were measured 24h after transfection in technical replicates. *Firefly* luciferase values are normalised to *Renilla*, independently expressed by the same vector and are shown as relative to that obtained for each construct cotransfected with the control miRNA mimic. Each dot represents and independent experiment. A construct containing three perfectly complementary binding sites for miR-125b (positive control) and the empty pmiRGlo vector (negative control) were included in the experiments. **C)** Gene Ontology enrichment analysis of the genes significanty up- and down-regulated upon miR-125b overexpression (T-RNA) and putative miR-125b direct targets (RIP-RNA/T-RNA > 1.5) performed with DAVID. The graph shows enrichment scores for one representative term for each cluster grouped by semantic similarities and including terms with a padj (Benjamini) < 0.05. See Supp Table 2 for a full list of terms. Error bars represent SEM. \**p* <0.05, ***p < 0.001, ****p<0.0001, paired Student t test values of the log[fold change] values.

**Table 1.**
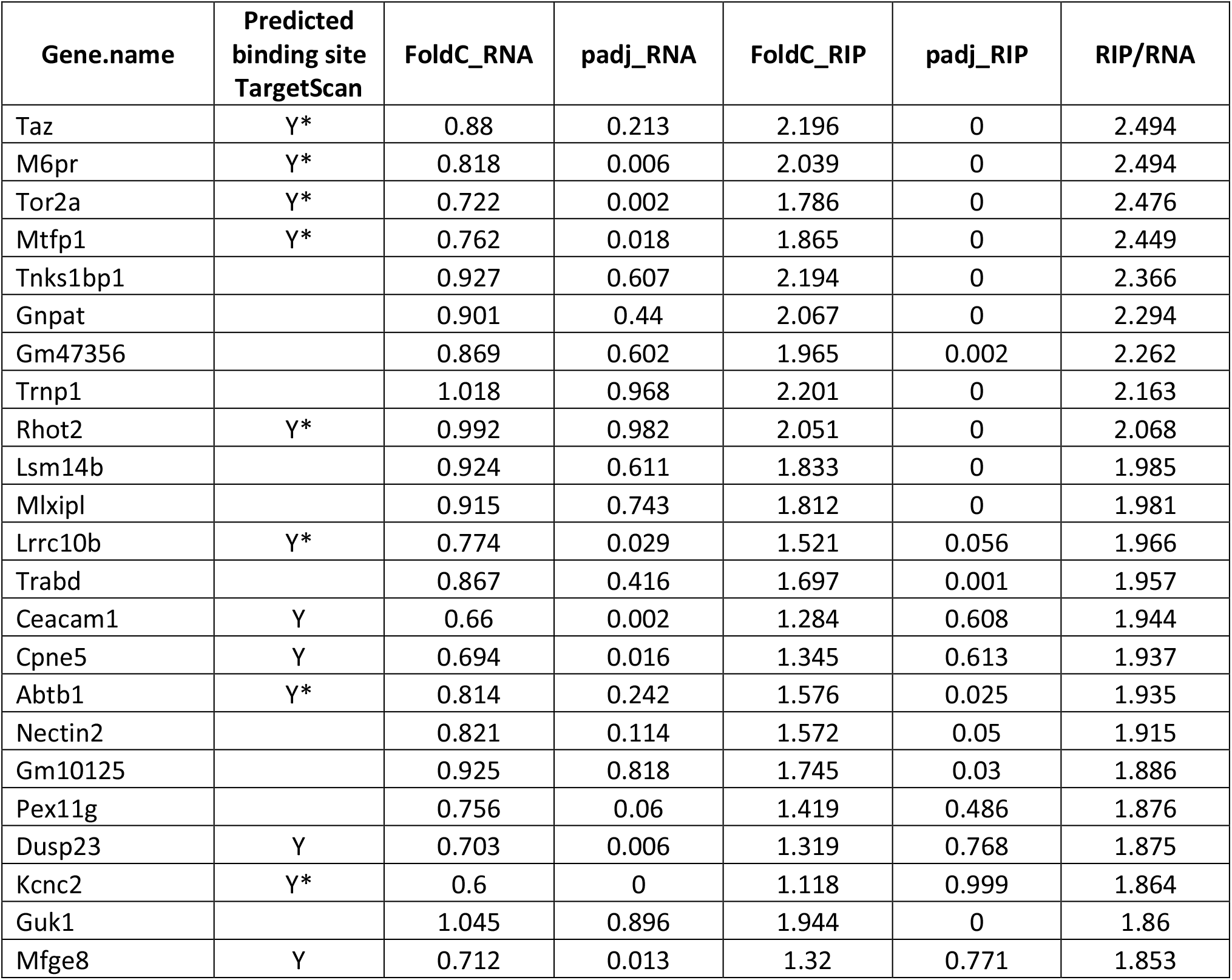
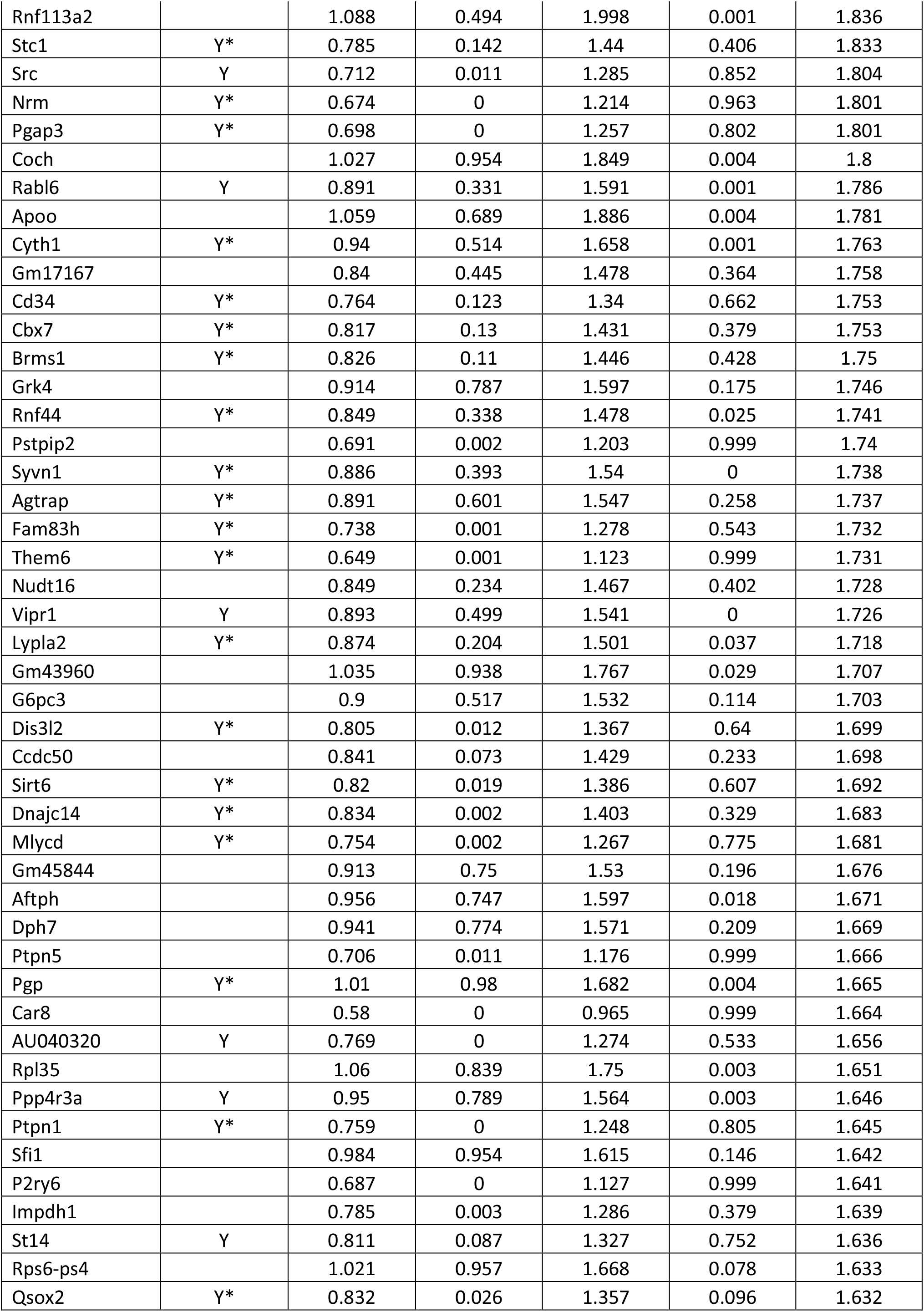
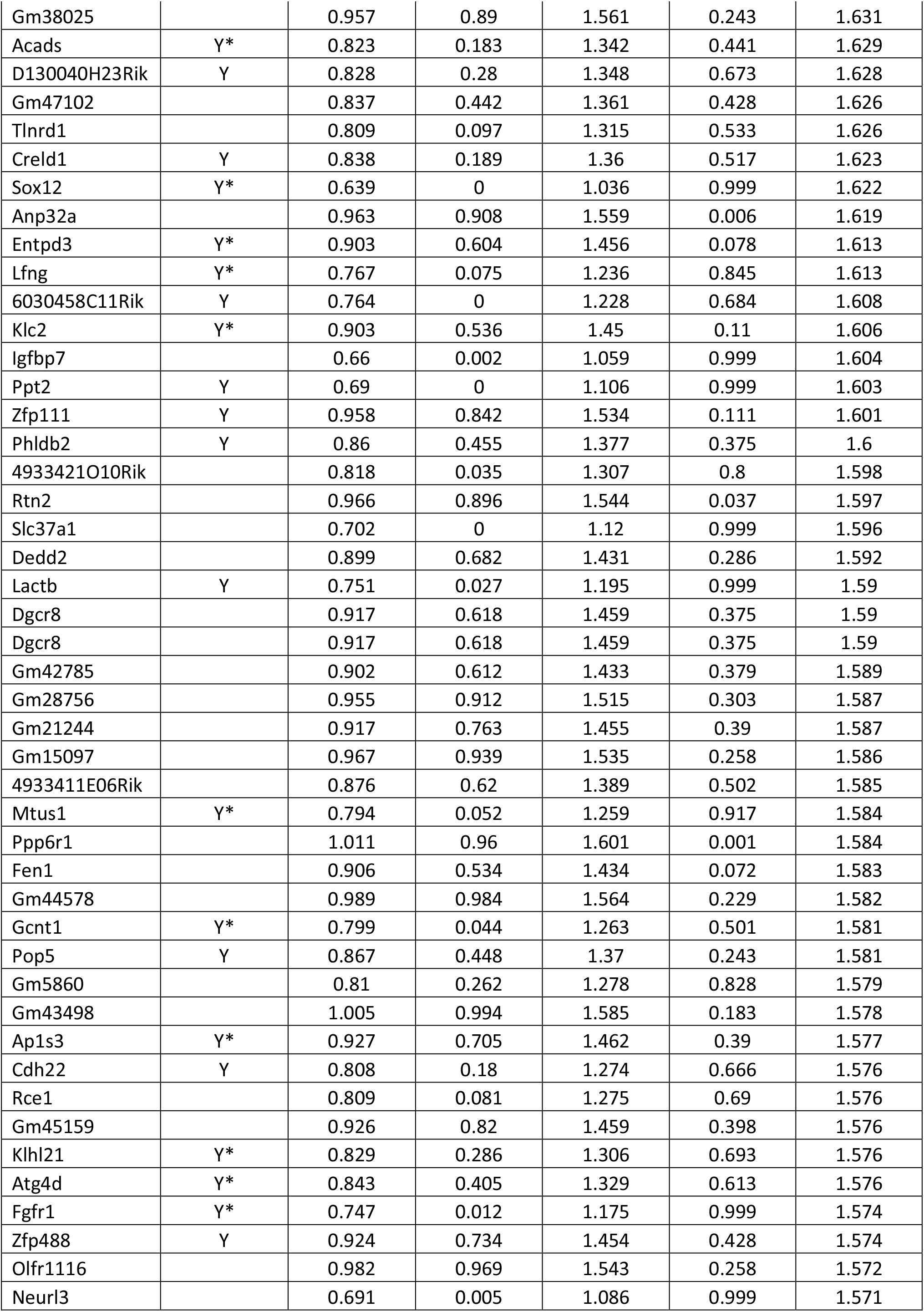
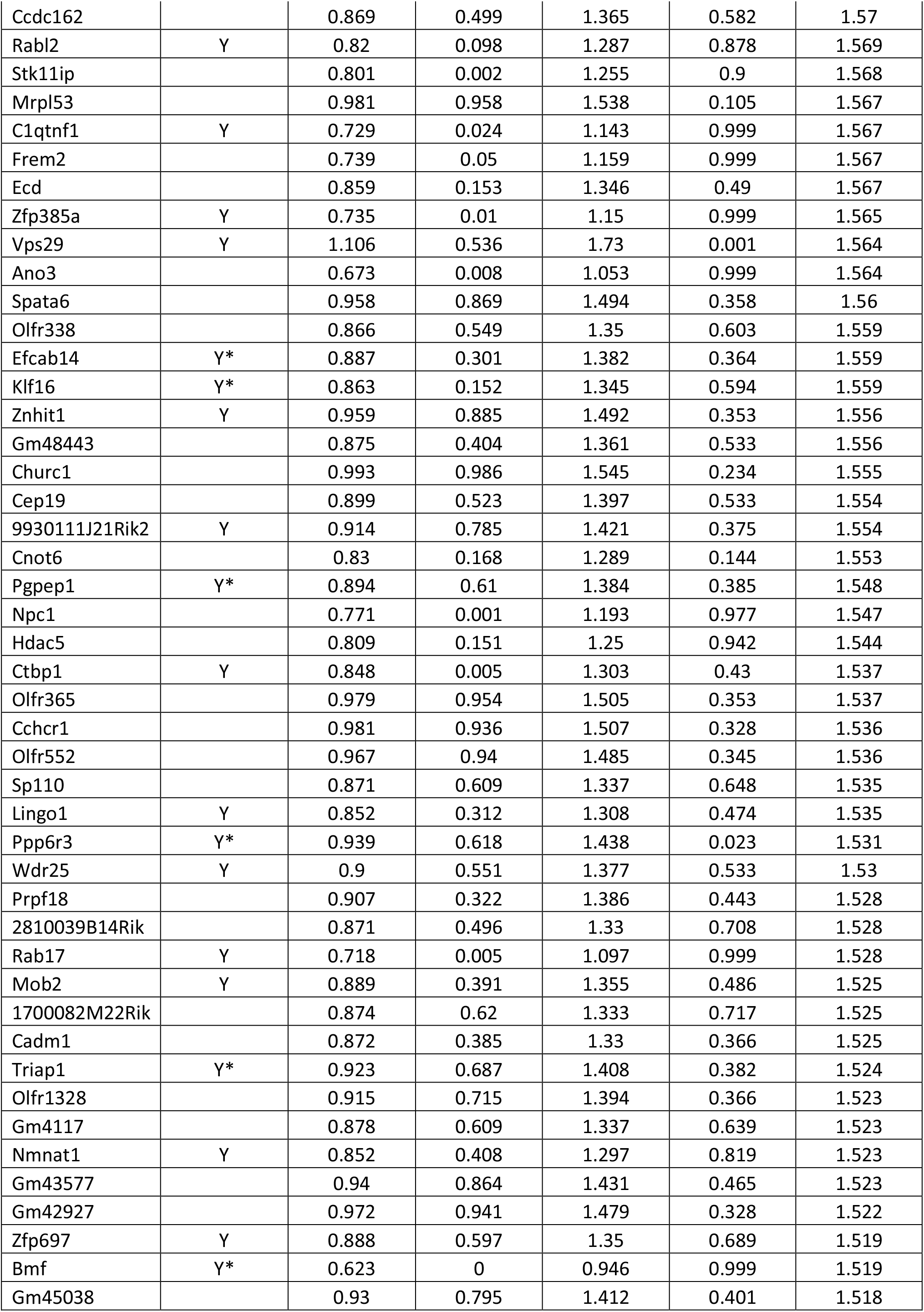
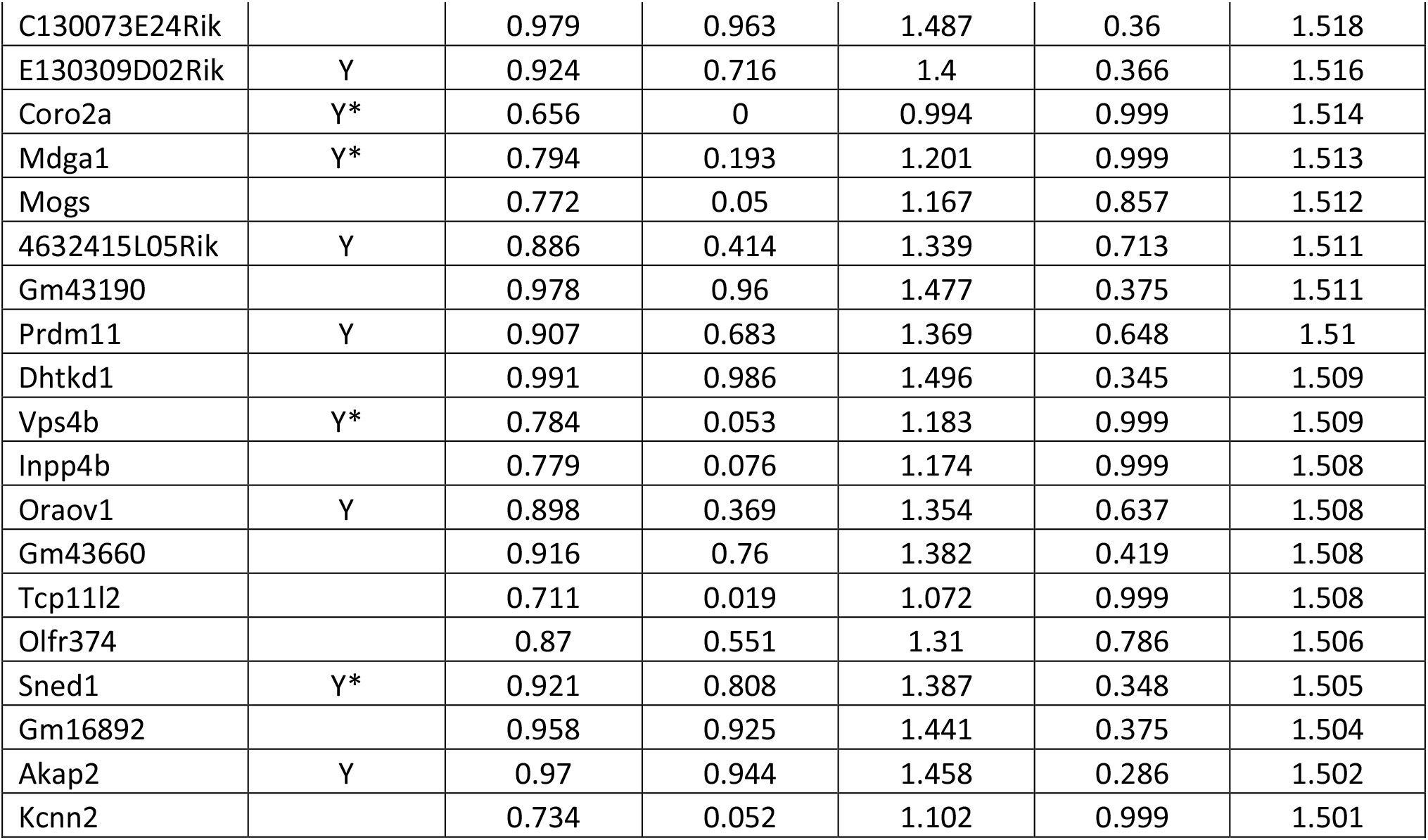
MiR-125b direct targets identified by integration of RIP-seq and T-RNA-seq differential analysis. Genes (180) with RIP-seq to T-RNA ratio > 1.5 are shown. FoldC_T-RNA and padj_T-RNA= Fold change and adjusted p-value of differential analysis of transcripts in MIN6 cells transfected with miR-125b mimics vs controls. FoldC_RIP and padj_RIP= Fold change and adjusted p-value of differential analysis of Ago2-immunoprecipitated RNAs in MIN6 cells transfected with miR-125b mimics vs controls. Predicted binding site TargetScan=Y indicates the presence of at least one predicted miR-125b binding site by TargetScan mouse v7.2. * (Y*) indicates target conservation in humans.

We further validated the six targets (*Taz, M6pr, Tor2a, Mtfp1, Tnks1bp1* and *Gnpat*) with the highest RIP-RNA/T-RNA ratios using luciferase reporter-based assays(20). As expected, co-transfection of miR-125b mimics reduced *Firefly/Renilla* activity ratios in cells transfected with constructs containing the *Taz*, *M6pr, Tor2a* and *Mtfp1* 3’UTRs, *Gnpat* CDS or three perfectly-complementary sequences to miR-125b (positive control), downstream the *Firefly* open reading frame (ORF) in the bicistronic plasmid pMiRGlo, demonstrating that miR-125b silence gene expression through those sequences (Fig. 2B). In contrast, miR-125b mimics did not affect *Firefly/Renilla* activity of the empty vector or in the presence of the 3’UTR of *Gnpat* which lacked predicted miR-125b binding sites. We failed to validate miR-125b action through *Tnks1bp1* CDS, perhaps due to the location of the binding site downstream the luciferase CDS and not within the CDS itself(22).

### MiR-125b targets genes encoding mitochondrial and lysosomal proteins

To identify biological pathways and functions regulated by miR-125b we submitted the list of genes dysregulated at the RNA level (T-RNA, 306 and 317 genes down- and up-regulated, respectively, padj < 0.1, Table S1) as well as those identified as miR-125b direct targets with a RIP/T-RNA >1.5 (Table 1) to a Database for Annotation, Visualization and Integrated Discovery, DAVID(23). Functional annotations with high enrichment scores included mitochondrion, tricarboxylate (TCA) cycle, sterol biosynthesis, protein transport, Golgi apparatus and endosome (Fig. 2C, Table S2), all closely associated to the capacity of the β-cells to produce and secrete insulin in response to glucose.

The gene at the top of our miR-125b-targets list, the lysosomal/Golgi-associated *M6pr* (Manose-6-phosphate receptor-cation dependent, M6PR-CD) controls trafficking of lysosomal hydrolases from the Golgi via endosomes(24) and thus may influence protein turnover and lysosomal function. Luciferase assays further confirmed the interaction of miR-125b *via* a predicted conserved miR-125b binding site in its 3’UTR (Fig. 3A) and Western-(Immuno) blot of EndoCβ-H1 cells overexpressing miR-125b and EndoCβH1-MIR125B2-KO cells showed a strong down- and up-regulation in M6PR protein, respectively (Fig. 3B) without significant changes at the mRNA level (Fig. 3C, Table S3). These results confirm that miR-125b targets M6PR to repress protein production in human EndoCβ-H1 cells..

**Figure 3.**
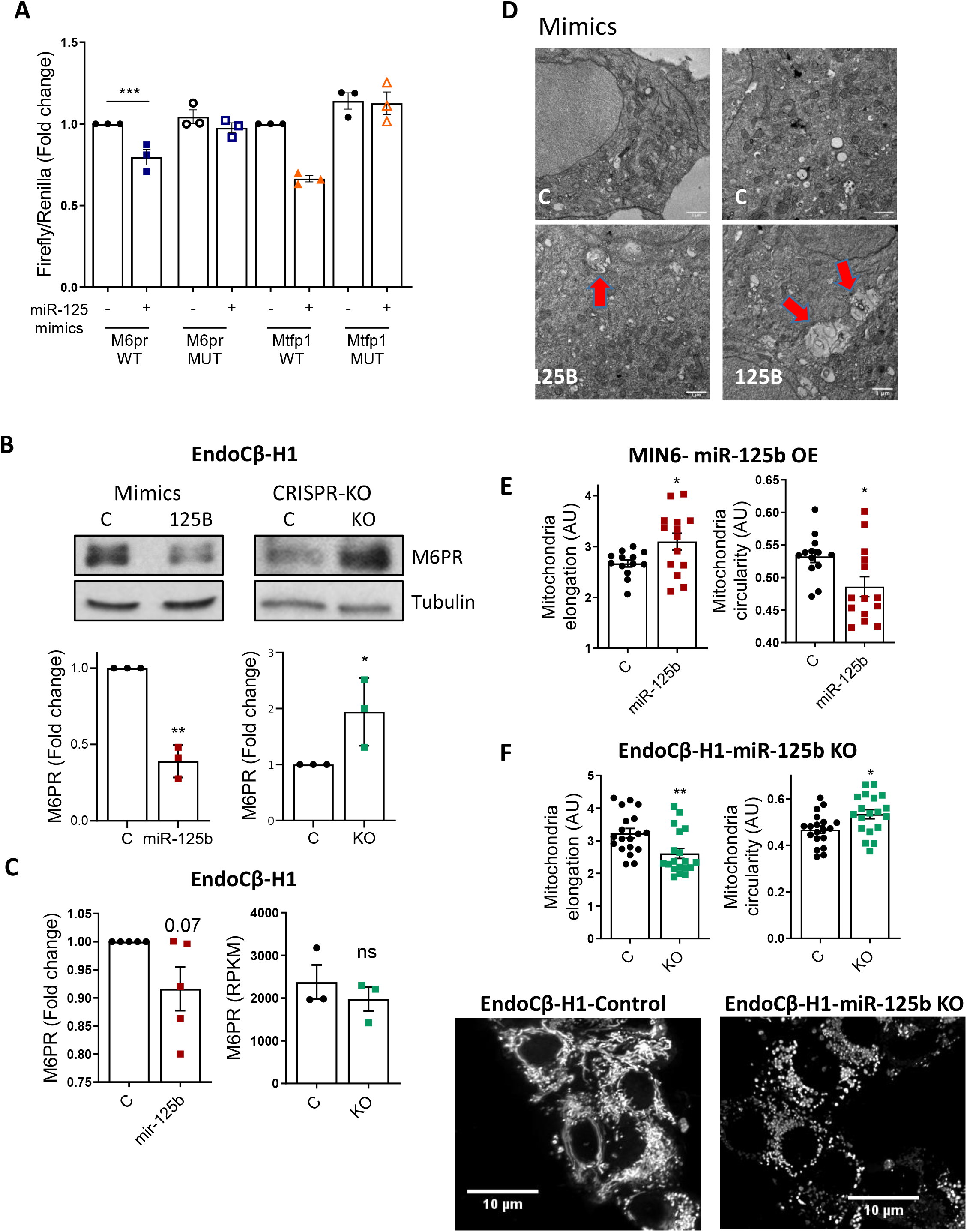
MiR-125b represses *M6pr* and *Mtfp1* and alters lysosomal and mitochondrial morphology. **A)** MIN6 cells were transfected with luciferase reporters containing the *M6pr* or the *Mtfp1* 3’UTRs with (mutated, MUT) or without (wild-type, WT) three/two point mutations in the sequence complementary to the miR-125b seed. Control (-) or miR-125b (+) mimics were co-transfected and *Firefly* luciferase values normalized to *Renilla*. Each dot represents and independent experiment. **B)** Representative Western blot showing reduced (left-hand side panel) and increased (right hand-side panel) M6PR protein levels upon transfection with 1nM miR-125b (miR-125b)/control (C) mimics or CRISPR-Cas9-mediated deletion of miR-125b in EndoCβ-H1 cells, respectively. Bar graphs show densitometry quantification of M6PR, using ImageJ, normalized by Tubulin and presented relative to the control. **C,** Left hand-side panel: RT-qPCR measurement of *M6PR* mRNA in EndoCβ-H1 cells transfected with control (C) or miR-125b (miR-125b) mimics, normalized by the housekeeping gene *Ppia* and presented as fold change of the control. Right hand-side panel: Normalized (Reads *per* Kilobase of transcript, *per* Million mapped reads (RPKM)) M6PR reads in Control *vs* MIR125B2-KO (KO) EndoCβ-H1 cells. **D,** Representative Transmission Electron Microscopy image of MIN6 cells transfected for 48h with control (C) or miR-125b (125B) mimics. Red arrows show enlarged lysosomal structures. Scale bar = 1 μm. These were not observed in any of the imaged controls (n=3 independent experiments). **E,F)** Quantitative analysis of mitochondria number and morphology on deconvoluted confocal images of (E) MIN6 cells transfected with miR-125b (red) or control (black) mimics or (F) EndoCβ-H1 with CRISPR/Cas9-mediated miR-125b deletion (green) or controls (black). Cells were stained with Mitotracker green. An ImageJ macro was generated and used to quantify individual mitochondria length (elongation) and circularity (0:elongated; 1:circular). Each dot represents one acquisition (n=3 (E), n=4 (F) independent experiments). Lower panels show representative confocal images of the mitochondrial network of EndoCβ-H1-MIR125B2-KO and control cells. Scale bar: 10 μm. Error bars represent SEM. \**p* <0.05, **p < 0.01, ***p<0.001, paired Student t test of the log[fold change] values (A-C); Welch t test (E, F).

We hypothesized that M6PR downregulation following miR-125b overexpression may lead to a defect in lysosomal hydrolases and an accumulation of lysosomal structures with non-digested cargos. Accordingly, electron microscopy (EM) revealed an accumulation of aberrant enlarged lysosomal structures in MIN6 cells overexpressing miR-125b (Fig. 3D).

As mentioned above, GO analysis of genes dysregulated upon miR-125b overexpression also revealed a significant over-representation of genes involved in mitochondrial function (Table S2, Fig. 2C), including one of the top miR-125b direct targets, mitochondrial fission process 1 (*Mtfp1*, also known as MTP18). Luciferase experiments further confirmed the interaction of miR-125b *via* a predicted conserved binding site in *Mtfp1* 3’UTR (Fig. 3A). MTFP1 has previously been implicated in mitochondrial fission and apoptosis in mammalian cells(25). To explore a possible role for miR-125b in controlling mitochondrial morphology, we stained MIN6 cells transfected with control or miR-125b mimics with a mitochondria-targeted green fluorescent probe (Mitotracker green). MIN6 cells overexpressing miR-125b contained the same number and overall mitochondrial area, though their mitochondria were slightly more elongated and less circular (Fig. 3E, S5A). In contrast, EndoCβH1-*MIR125B2-KO* cells revealed a marked reduction in mitochondrial area and mitochondria with a smaller perimeter, that were also less elongated and more circular (Fig. 3E, S5B). Further emphasizing the importance of miR-125b for mitochondrial homeostasis, GO analysis following RNAseq of EndoCβH1-MIR125B2-KO confirmed a strong enrichment in dysregulated genes associated with mitochondrial function (Fig. S5C, Table S4), including *MTFP1* (1.5 fold, p-value: 0.006, padj 0.1).

These findings suggest that relief from miR125-mediated repression leads to disruption of the tubule-vesicular network and to profound changes in the expression of genes involved in mitochondrial homeostasis which may underlie the observed improvement in insulin secretion.

### MiR-125b overexpression in β-cells impairs glucose tolerance *in vivo*

Given the regulatory effect of miR-125b *in vitro*, we decided to study the role of miR-125b *in vivo* by generating a mouse with β-cell specific, doxycycline-inducible overexpression of miR-125b using a RIP7-rtTA promoter (14). MIR125B-Tg (MIR125B-Tg: RIP7-rtTA^+/−^, MIR125B Tg^+/−^) animals contained ~60-fold more islet miR-125b than controls (Control: RIP7-rtTA^+/−^) (Fig. 4A), whereas we didn’t detect statistically significant changes in miR-125b levels in the hypothalamus, a potential site for activity of the RIP promoter (26) (Fig. S6A).

**Figure 4.**
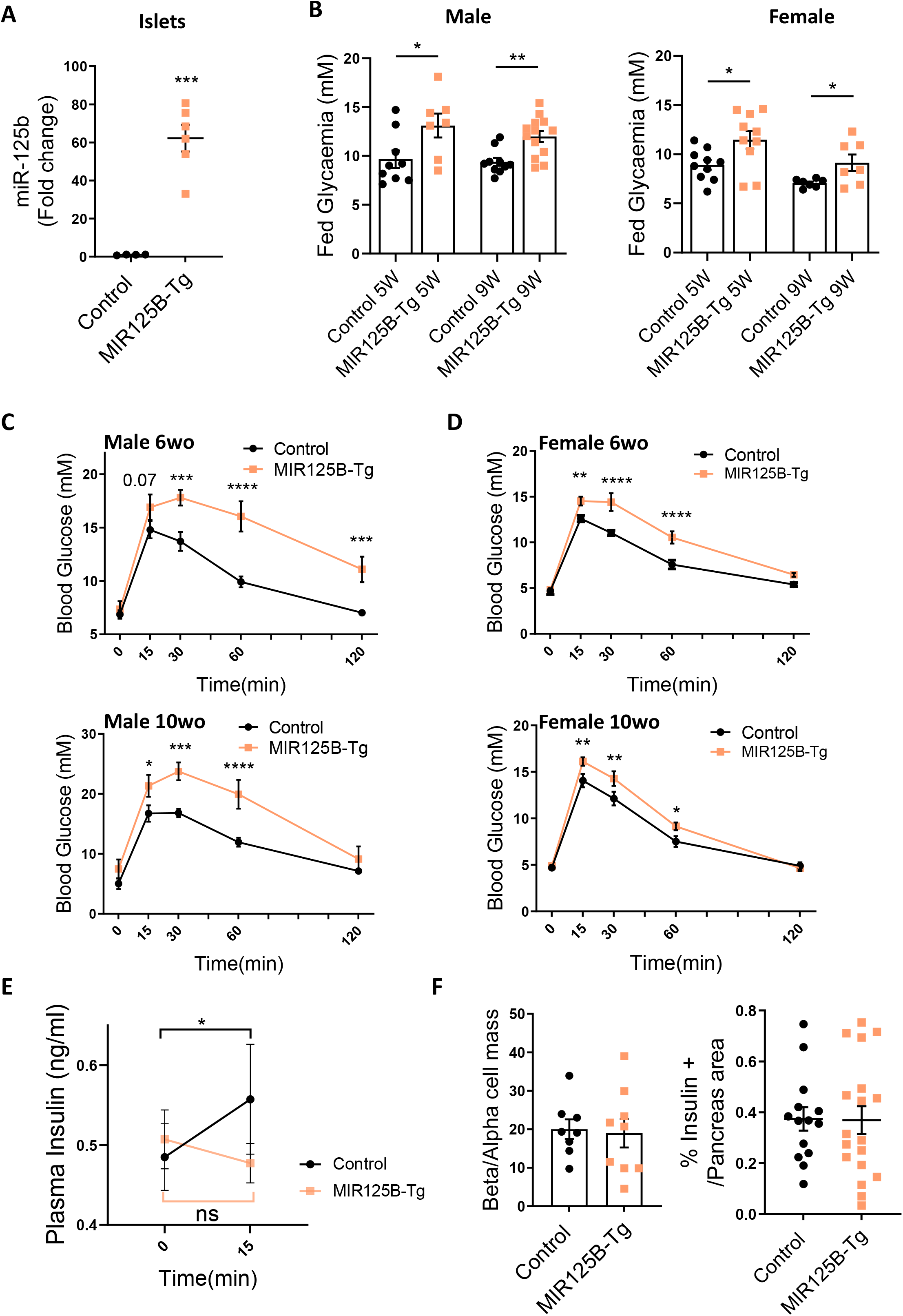
Mice with β-cell-selective overexpression of miR-125b are glucose intolerant and present impaired glucose stimulated insulin secretion. **A)** MiR-125b RT-qPCR in isolated islets from Control (RIP7-rtTA^+/−^) and MIR125B-Tg mice (RIP7-rtTA^+/−^, MIR125B Tg^+/−^). Data are fold change *vs* Control. **B)** Glycaemia in 5 and 9 week old Control and MiR125B-Tg mice fed *ad libitum*. For A and B, each dot represents a single mouse. **C, D)** Glucose tolerance test in 6 and 10 week old MIR125B-Tg and littermate control male (C) and female (D) mice. n=7-8C, 5-7Tg. **E)** Glucose (3 g/kg)-induced insulin secretion assessed in 10 week old MIR125B-Tg and littermate controls (n=13C, 11 Tg). **F)** Pancreata from MIR125B-Tg and littermate controls were fixed and subjected to immunocytochemical analysis for insulin and glucagon. β- and α-cell masses are presented as the β-cell/α-cell ratio and correspond to quantification of insulin-positive area/glucagon positive area ratio (left-hand side panel). β-Cell mass is presented as a percentage of the pancreatic surface and corresponds to quantification of the insulin-positive area per pancreas area quantified in whole pancreas sections. Each dot represents one pancreatic section with n = 3 mice/genotype. Error bars represent SEM. \**p* <0.05, **p<0.01, ***p < 0.001, ****p<0.0001, Welcht test (A, B, F); two-way ANOVA (repeated-measures), Fisher least significance different test (C-E).

MIR125B-Tg mice showed indistinguishable weight gain compared to controls (Fig. S6B) whereas random-fed glycaemia was significantly higher in MIR-125B-Tg in both males and females (Fig. 5B). Consistently, MIR125B-Tg mice were highly glucose intolerant (Fig. 4C,D), though fasting glycaemia was unaffected. No changes in glucose tolerance were observed in animals bearing the transgene in the absence of rtTA (-rtTA Control: RIP7-rtTA^-/-^, MIR125B OE^+/−^, Fig. S6C), demonstrating that the defects observed were not due to an off-target genomic event resulting from transgene integration. Blood glucose levels were efficiently reduced by administration of exogenous insulin excluding insulin insensitivity as a major player in the glucose intolerance observed in MIR125B-Tg animals (Fig. S6D). Moreover, plasma insulin did not increase in MIR125B-Tg mice in response to an intraperitoneal injection of glucose (Fig. 4E) suggesting impaired β-cell secretory function. β-cell mass and β- to α-cell ratio, as well as proliferation and apoptosis remained unchanged in the transgenic animals (Fig. 4F, Fig. S6E,F).

**Figure 5.**
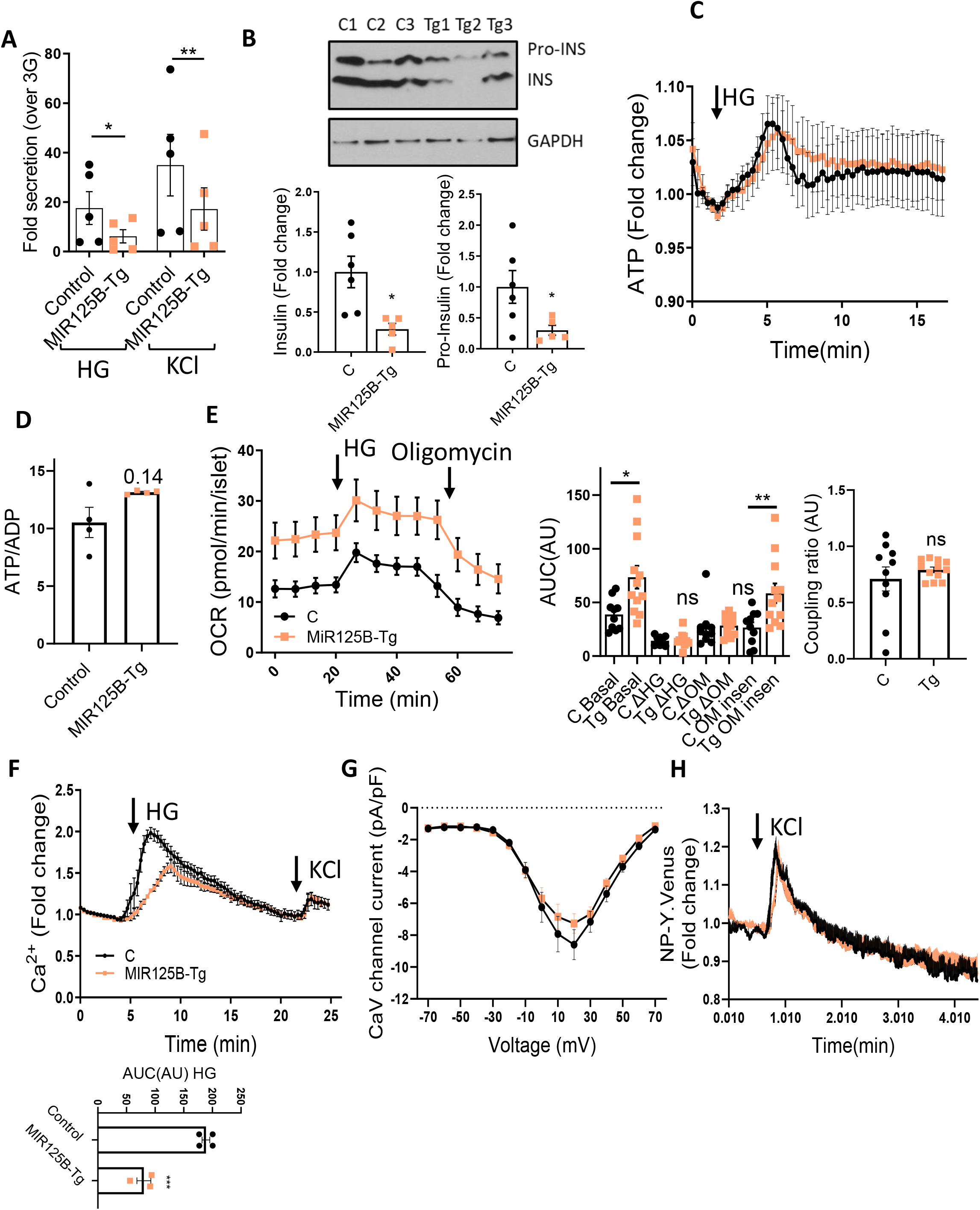
MIR125B-Tg islets contain and secrete less insulin than controls. **A)** Fold induction insulin secretion *vs* low glucose (3mM, LG) in response to 30 minutes high (17mM, HG) glucose or KCl (17mM KCl and 3mM Glucose) in islets from 10-11 week old Control (RIP7-rtTA^+/^-) and MIR125B-Tg (RIP7-rtTA^+/−^, MIR125B Tg^+/−^) mice. **B)** Insulin and pro-insulin content in 10 week old Control (C) and MIR125B-Tg (Tg) mice. A representative WB is shown with islets from 3 C and 3 Tg animals. Bar graphs show densitometry quantification of insulin and pro-insulin, using ImageJ, normalized by GAPDH and presented relative to the average control. **C)** ATP rises in response to high glucose (17mM *vs* 3mM) and **D)** ATP/ADP ratio at 3mM glucose in intact MIR125B-Tg and control islets infected with an adenoviral Perceval sensor. n=4-5 mice/genotype, with 1-2 acquisitions per mouse and an average of 5 islets/acquisition. **E)** Oxygen consumption rate (OCR) measured in the presence of 3mM or 17mG (HG) glucose and 5μM oligomycin (OM) over time as indicated. Islets were pre-incubated for 1h at 3mM glucose. Left hand-side panel shows traces from the actual experiment. Middle and right hand-side panels show calculations during the indicated periods. “Basal” shows AUC under basal (3mM) conditions, ΔHG and –ΔOM report the change in respiration when 17mM Glucose and oligomycin are added, respectively. The oligomycin insensitive value (OM insen) reports the residual respiration after the addition of oligomycin and coupling ratio (right-hand side panel) represents the oligomycin-sensitive respiration divided by respiration preceding oligomycin. Each dot represents a plate well with 3-6 size-matched islets (n=10-12) extracted from 3 Control (C) and 4 MIR125B-Tg (Tg) mice. **F)** Ca^2+^rises in response to high glucose (17mM *vs* 3mM) and KCl (20mM KCl, 3mM Glucose) in intact MIR125B-Tg and control islets incubated with Cal-520. n=4-5 mice/genotype, with 1-2 acquisitions per mouse and an average of 5 islets/acquisition. AUC corresponding to HG incubation was determined and presented on the bottom. **G)** Average maximum CaV current densities recorded from control and MIR125B-Tg β-cells in response to 10 mV steps from −70 to 70 mV. N= 16 and 29 β-cells from 2 control and 4 MIR125B-Tg mice. **H)** NP-Y.Venus fluorescence increases in response to 20mM KCl in cells from dissociated MIR125B-Tg and control islets infected with an adenoviral NP-Y-venus sensor. n=3-4C mice/genotype 2ith 2 acquisitions per mouse and 1-2 β-cells per acquisition. Each dot represents a single mouse in all graphs unless otherwise indicated. Error bars represent SEM. ns=not significant \**p* <0.05, **p<0.01, ***p < 0.001, two-way ANOVA (repeated-measures) and Bonferroni multiple comparisons test (A), paired Student (B) and unpaired Welch (D-E) or Student t test (F).

Taken together, these data suggest that increased miR-125b in the β-cell results in hyperglycaemia and glucose intolerance due to impaired β-cell secretory function.

### MiR-125b transgenic islets contain and secrete less insulin

To further explore a cell autonomous defect in insulin secretion, we measured GSIS in isolated islets from MIR125B-Tg mice which showed a strong reduction in insulin secretion in response to glucose or depolarisation by KCl (Fig. 5A, Fig. S6G). Consistent with our previous data in MIN6 cells, MIR125B-Tg islets contained less intracellular insulin (Fig. S5H), as further confirmed by Western blot (Fig. 5B, S6I), revealing lower levels of both intracellular pro-insulin and insulin. Accordingly, MiR125B-Tg pancreata contained less insulin (Fig. S6J). These data indicate a reduced capacity of MIR125B-Tg β-cells to produce and secrete insulin.

Upon entry in the β-cell, glucose is rapidly metabolized by mitochondria, causing a sharp increase in intracellular ATP/ADP ratio, closure of ATP-sensitive K^+^ (K_ATP_) channels, plasma membrane depolarization and Ca^2+^ influx into the cytosol to trigger insulin exocytosis(1). The fact that secretory responses to both high glucose and KCl were impaired in MIR125B-Tg islets suggested defects downstream of glucose metabolism. Accordingly, the glucose-mediated ATP/ADP rise and the basal ATP/ADP ratio were similar in MIR125B-Tg and Control isolated islets (Fig. 5C, D). OCR increases induced by high glucose and in response to the inhibitor of mitochondrial ATP synthase oligomycin were also comparable (Fig. 5E), confirming similar mitochondrial capacity to generate ATP. In contrast, both basal and oligomycin-insensitive respiration were significantly elevated in MIR125B-Tg islets (Fig. 5E), suggesting increased inner mitochondrial membrane proton conductance. Additionally, MIR125B-Tg islets showed a strong reduction in glucose-induced changes in intracellular Ca^2+^ (Fig. 5F). KCl-induced rises in Ca^2+^ were similar between MIR-125B-Tg and Control islets (Fig. 5F), suggesting that MIR125B-Tg β-cells contained functional voltage-gated calcium channels (VGCC). This was further confirmed in single β-cells by whole-cell voltage clamp that showed similar VDCC currents (Fig. 5G). The kinetics of individual fusion events were also identical in cells from MIR125B-Tg or Control islets (Figure 5H), as assessed by total internal reflection of fluorescence (TIRF) imaging of neuropeptide Y (NPY)-Venus-expressing vesicles near the plasma membrane(27), indicating that vesicle fusion itself was not affected by miR-125b overexpression.

### Enlarged lysosomes/autophagolysosomes are abundant in miR-125b transgenic islets

Our previous experiments in cell lines pointed to miR-125b as a regulator of mitochondrial and lysosomal function, which could contribute to defective insulin secretion. In contrast to our observations in cell lines, but consistent with an efficient capacity to generate ATP in response to glucose, mitochondrial area and morphology and the levels of mitochondrial DNA (mtDNA, Fig. 6B) were comparable in MIR125B-Tg and Control islets (Fig. 6A,B). Even though lysotracker staining revealed no differences in lysosomal area or numbers (Fig. S6K), transmission electron microscopy (TEM) identified the presence of enlarged lysosomes and autophagosomes in the MIR125B-Tg islets which were rarely found in control samples (Fig. 6C,D). Moreover, islets co-immunostaining identified a significant reduction in the lysosomal protease cathepsin D localized to (LAMP1-positive) lysosomes (Fig 6F). TEM also revealed a sharp reduction in the number of dense-core insulin granules, and an increase in granules with defective crystallization (light-core, abnormal rod-like or empty) (Fig. 6C,E). Importantly, fewer secretory granules were present within 200nm of the plasma membrane in MIR125B-Tg than in Control islets (Fig. 6E).

**Figure 6.**
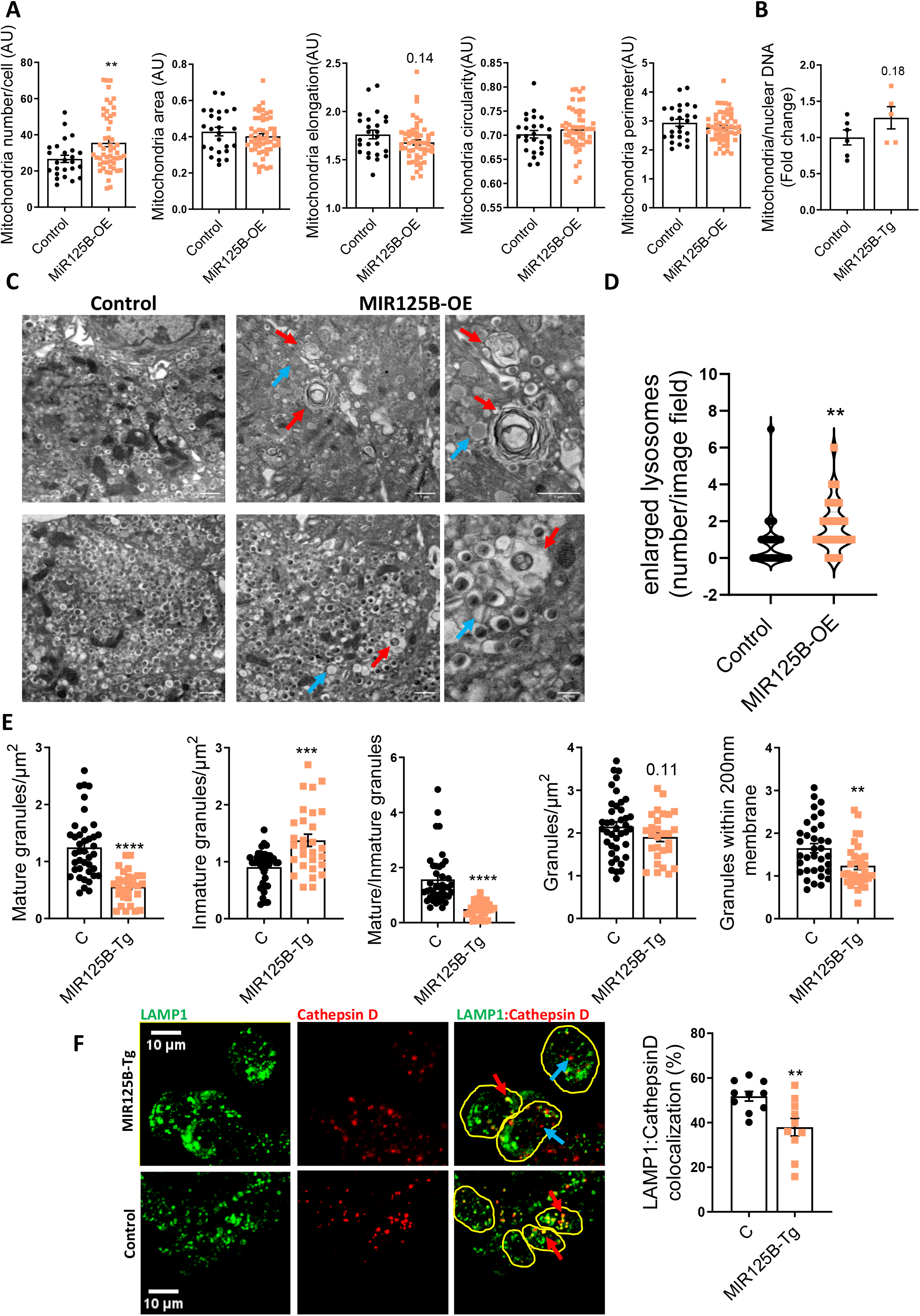
Enlarged lysosomes/autophagolysosomes are abundant in miR-125b transgenic islets. **A)** Quantitative analysis of mitochondria number and morphology on deconvoluted confocal images of dissociated MIR125B-Tg and Control islets from 10-11 week old mice. Cells were stained with Mitotracker green. An ImageJ macro was generated and used to quantify number of mitochondria per cell, total mitochondria area, individual mitochondria length (elongation), circularity (0:elongated; 1:circular) and perimeter. Each dot represents one cell (n=3 mice/genotype). **B)** Mitochondrial DNA copy number, calculated as the ratio of the mitochondrial encoded gene mt-Nd1 to the nuclear Cxcl12. Each dot represents a single mouse. **C)** Representative Transmission Electron Microscopy images of MIR125B-Tg and control β-cells. Red arrows: enlarged lysosomes with undigested cargos. Blue arrows: non-crystalized (“defective”) insulin granules, showing a rod-like structure or a light-core or empty interior. Scale bar = 1μm, 40nm (bottom right) interior. **D)** Number of enlarged lysosomes per field imaged. **E)**β-cell granule density. For (D, E) each dot represents one image (n=3 mice/genotype, 9-13 images/mouse). **F)** Representative confocal microscopy images of β-cells within MIR125B-Tg and control islets immunostained with anti-Cathepsin D (red) and anti-LAMP1 (green) antibodies. Yellow circles represent individual cells. Blue arrows show examples of only Cathepsin + particles and red arrows show particles with both LAMP1 and Cathepsin D + staining. ImageJ was used to quantify Cathepsin D/LAMP1 stained particles within each cell, represented in the right hand-side graph as percentage of LAMP1+ particles per cell. Each dot represents an individual islet with the average of 3-8 cells quantified per islet, extracted from 2 control (C) and 2 MIR125B-Tg mice. Error bars represent SEM. \**p* <0.05, **p<0.01, ****p < 0.0001, Welch t test.

### β-cell identity is altered in MIR125b-Tg islets

To shed more light into the molecular mechanisms underlying the secretory defects of miR-125B-Tg islets, we performed RNA-seq on MIR125B-Tg and Control islets. 320 and 398 genes were significantly (padj <0.1) down- and up-regulated, respectively (Table S5).

GO analysis of downregulated genes identified a strong enrichment within pathways and processes associated with mature β-cell function such as regulation of insulin secretion and endocrine pancreas development (Table S6, Fig. 7A,B, S7A), with a significant reduction of several important β-cell identity genes such as *Ucn3, Pdx1, NeuroD1, Nkx2.2, Slc2a2* and *Nkx6.1*. These genes don’t contain predicted miR-125b binding sites or had been identified as direct miR-125b targets in our RIP-seq experiments, suggesting an indirect effect. Further supporting a loss of β-cell identity, up-regulated genes were significantly associated with neuronal features (Table S6, Fig. 7A,S7B). We also observed an upregulation of many glycoprotein genes and of genes associated with cellular and focal adhesion, signalling pathways and voltage-gated and potassium channels. Golgi-associated genes were both up and down-regulated Table S6, Fig. 7A, S7B).

**Figure 7.**
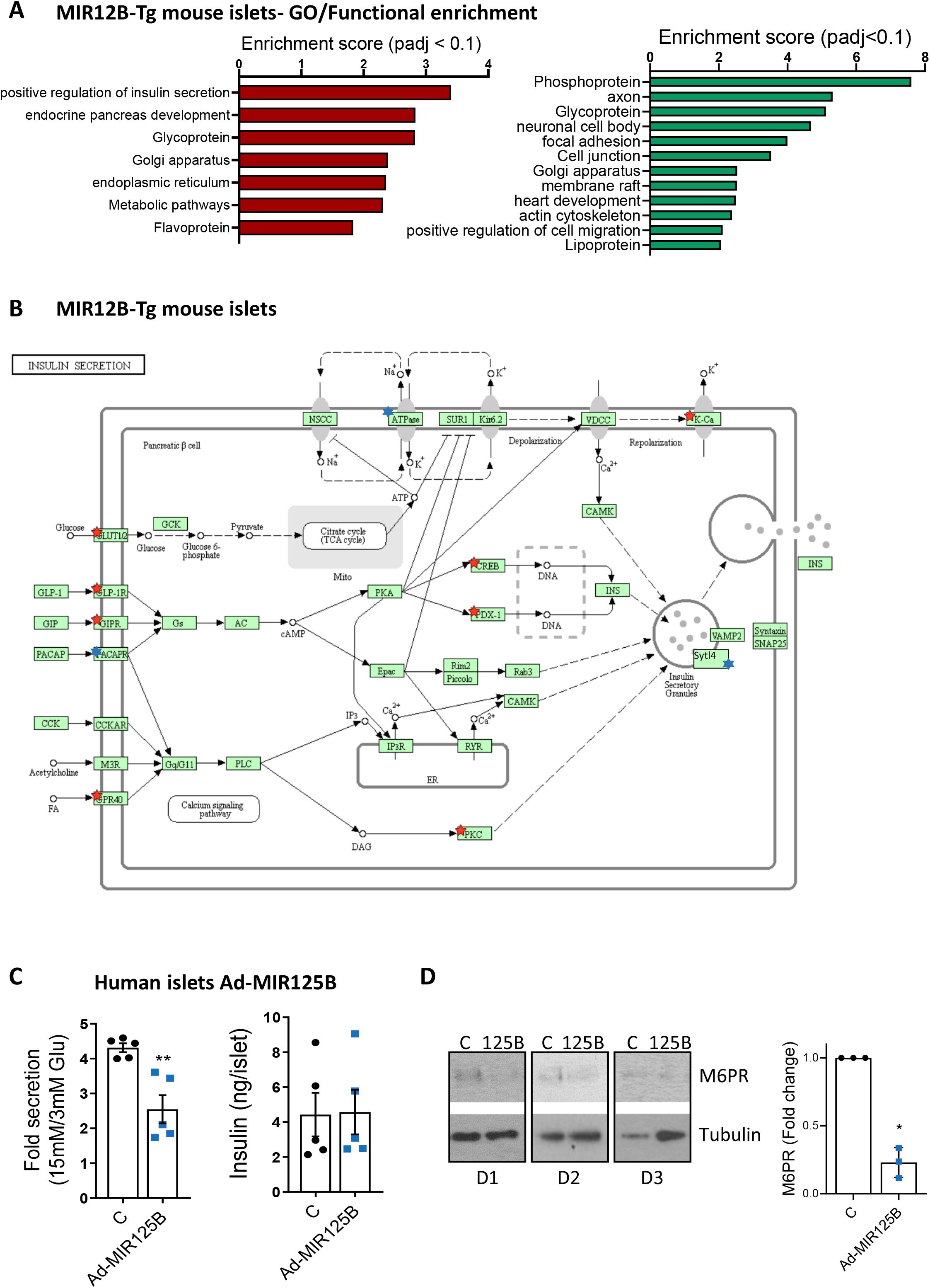
MiR125b overexpression alters gene expression in mouse and human islets. **A)** Gene Ontology analysis of significantly (padj <0.1) down-regulated (left, red) and up-regulated genes (right, green) in MIR125B-Tg islets from 6-week old mice *vs* controls performed with DAVID. The graph shows enrichment scores for one representative term for each cluster grouped by semantic similarities and including terms with padj (Benjamini) < 0.05. See Supp Table 6 for a full list of terms. **B)** KEGG pathway “Insulin secretion”. Red and blue stars indicate significantly down- and up-regulated genes in MIR125B-Tg islets *vs* control, respectively. *Sytl4* (granuphilin) has been manually added to the pathway. **C)** Glucose-stimulated insulin secretion (left hand-side panel) and insulin content (right hand-side panel) quantified after 30 minutes of 15 mM glucose stimulation following 1h pre-incubation at 3 mM glucose of human islets infected with adenovirus expressing miR-125b (Ad-MIR125B) or a non-targeting control (C) at 5 MOI 48h before the experiments. GSIS gata is presented as fold change of basal level. Each dot represents an independent experiment performed with islets from two different donors. **D)** Western blot showing reduced M6PR protein levels infection of dissociated human islets from three different donors (D1-3) infected with Ad-MIR125B or control adenovirus (2 MOI) for 48h. Bar graphs show densitometry quantification of M6PR, using ImageJ, normalized by Tubulin and presented relative to the control.

### MiR-125b alters insulin secretion and gene expression in human islets

To further explore the relevance of miR-125b in human β-cells, we performed GSIS assays in human islets infected with Ad-MIR125B, achieving a ~4.5 fold increase in miR-125b levels (Fig. S8A). MiR-125b overexpressing islets secreted ~50% less insulin than controls, though their insulin content remained unchanged (Fig. 7C, S8B). Additionally, we performed RNA-seq in dissociated human islets infected with these virus, achieving a ~3 fold increase in miR-125b (Fig. S8C). Principal component analysis (PCA) showed that most of the variation was due to the donor origin of the islets (Fig. S8D) and, thus this experiment was underpowered (~20% to detect a 2x Fold change with p<0.01, calculated using Scotty(28)). Nevertheless the expression of hundreds of genes tended to be altered (p-value <0.05, Table S7) and Gene Set Enrichment analysis of all the genes ranked by gene expression fold change identified a significant enrichment in several biological pathways (Fig. S8E,F, Table S8) notably including lysosomal (p<0.0001, FDR=0.07) and calcium signalling (p<0.0001, FDR=0.06). Further suggesting a role for miR-125b in lysosomal function, a downregulation in M6PR protein was observed in dissociated human islets infected with Ad-MIR125B (Fig. 7D).

## DISCUSSION

Even though elevated levels of circulating miR-125b in association with higher HbA1c in T1D(29) and T2D(30) have been previously identified, this is the first study to demonstrate that miR-125b expression is induced by glucose in mouse and human islets in an AMPK-dependent manner. Although our capacity to demonstrate a correlation between hyperglycaemia and islet miR-125b levels in human islets *in vivo* was limited by the lack of HbA1c data, we found a significantly positive correlation with donor BMI which often associates itself with HbA1c(31).

Despite being highly expressed in β-cells, the function and mechanism of action of miR-125b in these cells have remained elusive until now. Here, we show for the first time that elevated expression of this miRNA in β-cells impairs their secretory function *in vitro* and *in vivo*.

We have generated a novel transgenic model capable of β-cell selective overexpression of miR-125b. These animals were hyperglycaemic and strongly glucose intolerant and presented a drastic reduction in circulating insulin following a glucose challenge.

Our findings suggest two main causes for the secretory defects in the MIR125b-Tg mice: strong impairment in GSIS and reduced insulin content. Unlike ATP, cytosolic Ca^2+^ increases following glucose stimulation were substantially lower in MIR125b-Tg islets though, paradoxically, similar levels of Ca^2+^ were observed upon membrane depolarization with KCl, even though KCl-stimulated insulin secretion was also strongly impaired. This points towards defects at different levels on the secretory pathway, as supported by the striking changes in gene expression. First, K_ATP_-independent amplification pathways(32) might be impaired in MIR125B-Tg islets, which display lower levels of genes encoding enzymes limiting for the production of guanine nucleotides and glutamine such as *Imphd1* and *GLUL* that can promote insulin exocytosis in an ATP/ADP independent manner(33; 34). Secondly, voltage-clamp electrophysiology experiments confirmed the presence of functional voltage-dependent Ca^2+^-channels in MIR125B-Tg β-cells. MIR125B-Tg islets expressed normal levels of Ca^2+^-channels but higher levels of PMCA3 and the Na^+^/Ca^2+^-exchanger NCX3, members of protein families responsible for Ca^2+^ extrusion in β-cells, which could potentially contribute to reduced levels of cytosolic calcium. Thirdly, TEM revealed fewer granules in close proximity to the plasma membrane in MIR125B-Tg β-cells and our RNAseq identified a strong alteration in genes involved in cellular granule docking and fusion such as granuphilin (*Sytl4*), the SNARE protein *Vamp3* and synaptotagmins (*Syt1/12/17*) (35; 36), pointing towards defects in the final steps of exocytosis. Finally, MIR125B-Tg islets present defective granule crystallization which might be partially due to reduced levels of the zinc transporter *Znt8* mRNA (37). Loss of β-cell identity might further contribute to these secretory defects, though some of these transcriptional changes may occur as a consequence of the moderate fed hyperglycaemia in these animals(38) or be residual from impaired maturation since RIP7-rtTA-driven transgene expression is expected to occur from E11,5.

Here, we unbiasedly identified dozens of miR-125b targets through the integration of RIP-seq and RNA-seq data(8; 20) of miR-125b overexpressing MIN6 cells. Given the supra-physiological levels of miR-125b achieved in these experiments, additional loss-of-function experiments will be key to confirm the targeting. It remains to be studied whether some of these targets are shared with the family member miR-125a, also expressed in β-cells. Additionally, some of these targets might not be conserved in humans and this may explain the fact that miR-125b modulation affects insulin content in mouse β-cells but less so in humans. Intriguingly, dishevelled binding antagonist of beta catenin 1 (*Dact1*) was not within this list and remained unchanged upon miR-125 manipulation in all our experimental models. *Dact1* was previously identified as a miR-125b target in β-cells, perhaps due to the presence of other pancreatic, highly proliferative cells in the samples(39)..

One of the genes at the top of our list of miR-125b targets, *M6pr*, encodes the cation-dependent manose-6-phosphate protein MP6R (CD-M6PR). Even though the function of CD-M6PR remains unknown in β-cells, this receptor is necessary for adequate lysosomal targeting of several hydrolases such as Cathepsin D(40).Remarkably, MIR125B-Tg islets contained less cathepsin D in their lysosomes, which appeared enlarged similarly to those observed by Masini et al. (41) in T2D islet micrographs, suggesting the accumulation of undigested cargos. Defects on the lysosomal/trafficking machinery can also affect the maturation, packaging and location of the secretory granules within the cell(42) and might contribute to the reduced insulin content and crystallization defects observed in the MIR125B-Tg mice. Whether elevated miR-125b during hyperglycamia/loss of AMPK activity contribute to these processes in T2D and how, remains to be studied.

Even though we had identified several genes encoding mitochondrial proteins as miR-125b targets in MIN6 cells, no mitochondrial morphological defects were detected in the transgenic islets, suggesting that endogenous miR125b levels are sufficient to exert maximal effects on mitochondrial structure. However, several mitochondrial genes were dysregulated in MIR125B-Tg islets and seahorse experiments showed a significant increase in basal and oligomycin-insensitive respiration, suggesting certain extent of mitochondrial dysfunction. On the other hand, deletion of miR-125b in EndoCβ-H1 cells resulted in a strong alteration of mitochondrial morphology and in the expression of mitochondrial proteins, including the miR-125b targets *Mtfp1* and *Gnpat. Mtfp1* is targeted by miR-125b in monocytes to promote elongation of the mitochondrial network (43). GNPAT acts to stabilize DRP1(44), which is required for mitochondrial fission and which deletion impaired insulin secretion(45; 46). In line with these earlier studies, EndoCβ-H1-MIR125B2-KO cells contained shorter mitochondria which correlated with increased GSIS. It has, however, been contested whether mitochondrial fragmentation promotes apoptosis or correlates with hyperglycaemia or diabetes(15; 47). Importantly, we did not observe any differences in cellular death or number in EndoCβ-H1-MIR125B2-KO neither in *Ddit3* (CHOP) mRNA levels, suggesting an absence of mitochondrial-induced ER stress in these cells. Future experiments are required to determine whether the positive effects of miR-125b elimination in β-cell function persist under conditions of cellular stress.

In summary, our results suggest that β-cell miR-125b has the potential to act as a glucose-regulated metabolic switch between the lysosomal system and mitochondria dynamics. Whole-body *MIR125B-2* knockout mice on a high-fat diet develop insulin resistance and glucose intolerance possibly due to the white fat accumulation (48). Nevertheless, insulin secretion or an effect in other metabolic organs were not assessed and our study provides compelling evidence of a role for miR-125b in controlling insulin secretion and strongly indicates that non-targeted administration of miR-125b mimics may reach the islet and lead to defective β-cell function and worsen diabetes.

Future studies focussed on β-cell-specific inhibition of miR-125b and on the function of specific targets will be essential to confirm a potential therapeutic benefit of targeting this miRNA or its targets for the treatment of diabetes.

## AUTHOR CONTRIBUTIONS

R. Cheung & G. Pizza (equal contribution) performed most experiments, designed research studies and obtained and analyzed data; P. Chabosseau performed research studies (TIRF) and designed microscopy macros. D. Rolando assisted with computational analysis of RNA-seq and RIP-seq data. A. Tomas and T. Burgoyne performed TEM experiments. Z. Wu, A. Salowka, A. Thapa, A. Macklin and Y. Cao performed research studies (insulin secretion, β-cell mass and apoptosis and Western blot, respectively). M.-S. Nguyen-Tu assisted with animal studies (*In vivo* Insulin secretion). M. T. Dickerson and David Aaron Jacobson performed whole-cell voltage clamp experiments. P. Marchetti, K. Bouzakri, J.Shapiro, L. Piemonti and E. de Koning, provided human islets. I. Leclerc assisted with maintenance of animal colonies. K. Sakamoto and D. M. Smith provided C13 and C991, respectively. G. A. Rutter contributed to the study design, with reagents and AMPK and LKB1 KO animals, and in writing the manuscript. A. Martinez-Sanchez conceived the study, designed and performed experiments, analyzed data, and wrote the paper; All authors read and approved the manuscript. A. Martinez-Sanchez is the guarantor of this work and, as such, had full access to all the data in the study and takes responsibility for the integrity of the data and the accuracy of the data analysis.

## ACKNOLEDGEMENTS

The authors thank Stephen M. Rothery from the Facility for Imaging by Light Microscopy (FILM) at Imperial College London for support with confocal and widefield microscopy image acquisition and analysis.

A.M-S was supported by an MRC New Investigator Research Grant MR/P023223/1. G.A.R. was supported by a Wellcome Trust Senior Investigator Award (WT098424AIA) and Investigator Award (212625/Z/18/Z) and MRC Programme grants (MR/R022259/1,MR/J0003042/1, MR/L020149/1). A.T was funded by the MRC (MR/R010676/1). IL was supported by a Diabetes UK project grant (16/0005485). This project has received funding from the European Union’s Horizon 2020 research and innovation programme via the Innovative Medicines Initiative 2 Joint Undertaking under grant agreement No. 115881 (RHAPSODY) to G.R. and P.M. Human islet preparations (Milan, Italy) were obtained from the European Consortium for Islet Transplantation; the Human Islet Distribution program was supported by Juvenile Diabetes Research Foundation Grant 3-RSC-2016-160-I-X.

## PRIOR PRESENTATION

Parts of this study were presented at the 79th Scientific Sessions of the American Diabetes Association, San Francisco, CA, 7–11 June 2019.

