## Supplementary material for "Glucose-dependent miR-125b is a negative regulator of β-cell function": Sup_Table9

| **Supplemental Table 9.** **Donor characteristics of human islets as provided by the isolation center**. | | | | |
| --- | --- | --- | --- | --- |
| **Identifier** | **BMI (Kg/m^2^)** | **Age** | **Sex** | **Origin (Facility)** |
| P43 | 27.4 | 72 | Male | Pisa, Italy |
| P59 | 21.5 | 68 | Male | Pisa, Italy |
| P60 | 27.8 | 61 | Male | Milano, Italy |
| P66 | 31 | 52 | Male | Pisa, Italy |
| P74 | 24.5 | 83 | Male | Pisa, Italy |
| P76 | 22.9 | 75 | Male | Pisa, Italy |
| P80 | 35 | 54 | Male | Edmonton, Cananda (McDonald) |
| P81 | 29.7 | 65 | Male | Edmonton, Cananda (McDonald) |
| P84 | NS | 89 | Male | Pisa, Italy |
| P86 | 23 | 41 | Male | Leiden, Netherlands |
| P89 | 24.5 | 64 | Male | Pisa, Italy |
| P94 | 29.4 | 51 | Male | Edmonton, Cananda (McDonald) |
| P95 | 42.60 | 38 | Male | Edmonton, Cananda (McDonald) |
| P97 | 28.7 | 62 | Male | Edmonton, Cananda (Shapiro) |
| P100 | 32.6 | 42 | Male | Pisa, Italy |
| P112 | 28.8 | 69 | Male | Edmonton, Cananda (McDonald) |
| P115 | 30.5 | 27 | Male | Edmonton, Cananda (Shapiro) |
| P118 | 28.4 | 68 | Male | Pisa,Italy |
| P119 | 29.4 | 49 | Male | Edmonton, Cananda (Shapiro) |
| P124 | 22.9 | 18 | Male | Edmonton, Cananda (Shapiro) |
| P125 | 24.5 | 75 | Male | Pisa, Italy |
| P128 | 22.5 | 74 | Male | Pisa, Italy |
| P119 | 29.4 | 49 | Male | Edmonton, Cananda (Shapiro) |
| P130 | 23.2 | 84 | Male | Pisa, Italy |
| P131 | 32 | 56 | Male | Milano, Italy |
| P132 | 24 | 57 | Male | Oxford, UK |
| P136 | 19.8 | 59 | Male | Pisa, Italy |
| P138 | 21.6 | 63 | Male | Milano, Italy |
| P140 | 23 | 41 | Male | Leiden, Netherlands |
| P141 | 31 | 45 | Male | Leiden, Netherlands |
| P142 | 31.2 | 71 | Female | Pisa, Italy |
| P148 | 26.4 | 66 | Male | Pisa, Italy |
| P152 | 33.8 | 44 | Male | Edmonton, Cananda (McDonald) |
| P28 | NS | 69 | Female | Edmonton, Cananda (Shapiro) |
| P36 | 25.2 | 37 | Female | Geneva, Switzerland |
| P41 | 17.9 | 58 | Female | Edmonton, Canada (Shapiro) |
| P42 | 21 | 23 | Female | Edmonton, Canada (Shapiro) |
| P77 | 33.1 | 66 | Female | Edmonton, Cananda (McDonald) |
| P85 | 23.9 | 62 | Female | Pisa, Italy |
| P87 | 23 | 66 | Female | Pisa, Italy |
| P93 | 26 | 57 | Female | Milano, Italy |
| P102 | 31.5 | 78 | Female | Edmonton, Cananda (Shapiro) |
| P106 | 20.6 | 49 | Female | Milano, Italy |
| P113 | 32.3 | 30 | Female | Edmonton, Cananda (McDonald) |
| P114 | 35 | 46 | Female | Oxford, UK |
| P116 | 26 | 55 | Female | Milano, Italy |
| P117 | 24.5 | 52 | Female | Milano, Italy |
| P120 | 25.8 | 58 | Female | Pisa, Italy |
| P123 | 22 | 88 | Female | Pisa, Italy |
| P126 | 18.5 | 66 | Female | Edmonton, Cananda (McDonald) |
| P134 | 44.1 | 51 | Female | Geneva, Switzerland |
| P137 | 28.2 | 84 | Female | Pisa, Italy |
| P142 | 31.2 | 71 | Female | Pisa, Italy |
| P144 | 31.2 | 49 | Female | Pisa, Italy |
| P145 | 25.7 | 78 | Female | Pisa, Italy |
| P146 | 31.9 | 73 | Female | Strasbourg, France |

Donors used for C991/C13 experiments (Figure 1E): P28, P36, P41, P42, P43

Donors used for Ad-AMPK-DN experiments (Figure 1E): P41, P43, P144 and P146

Donors used for effect of glucose on miR-125b expression (Figure 1A): P59, P80, P85, P95, P106, P114 and P116

Donors used for GSIS (Figure 1C): P144 and P146

Donors used for analysis of M6PR expression: P144, P145 and P146.

Donors used for RNAseq (Figure S8): P130, P132 and P138

All other samples used for correlation of miR-125b with BMI/Age/Sex (Figure 1b, Sup Figure 1)
