## Supplemental Figures for "Glucose-dependent miR-125b is a negative regulator of β-cell function"

**Supplemental Figure 1**

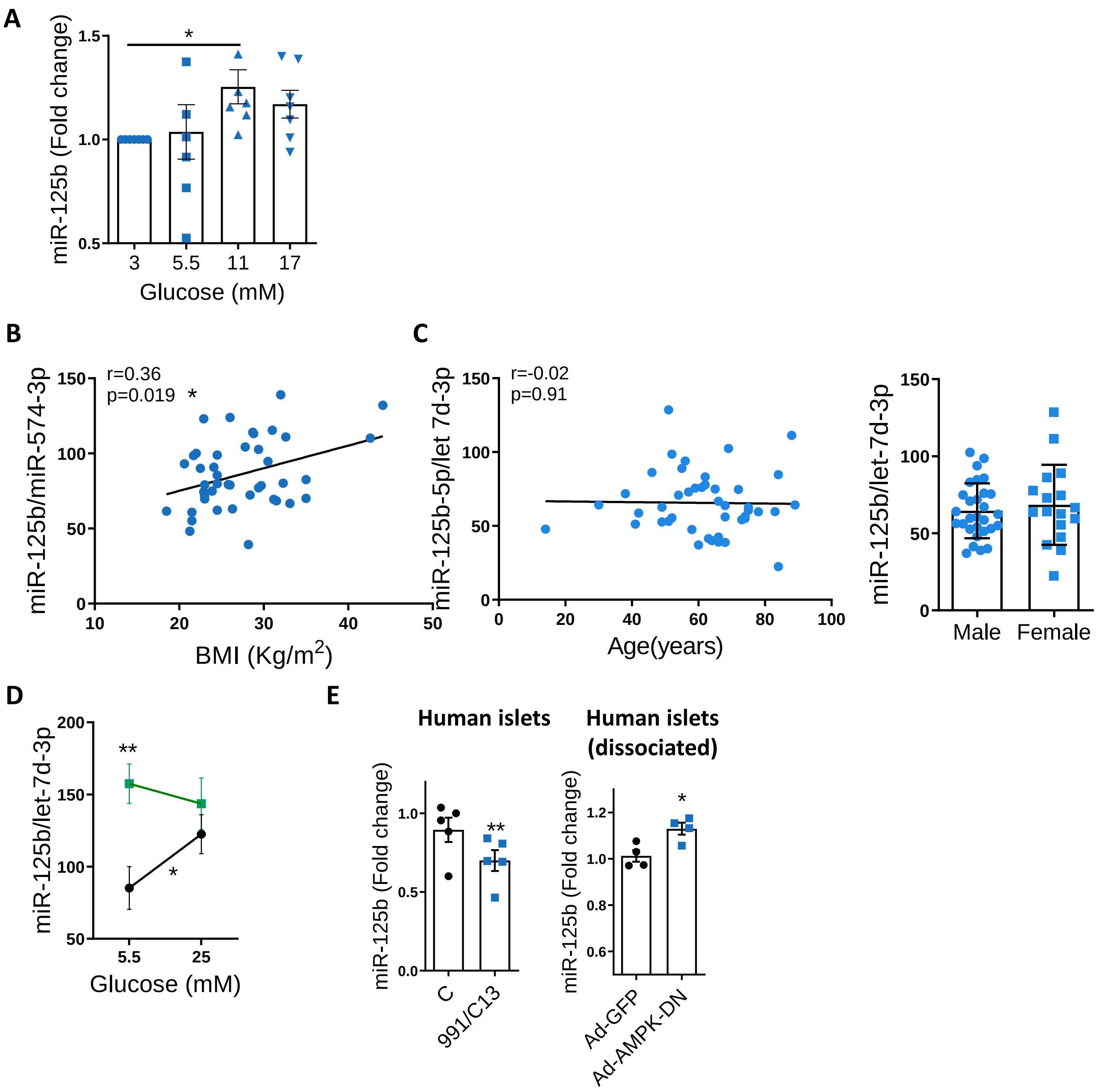

**Supplemental Figure 2**

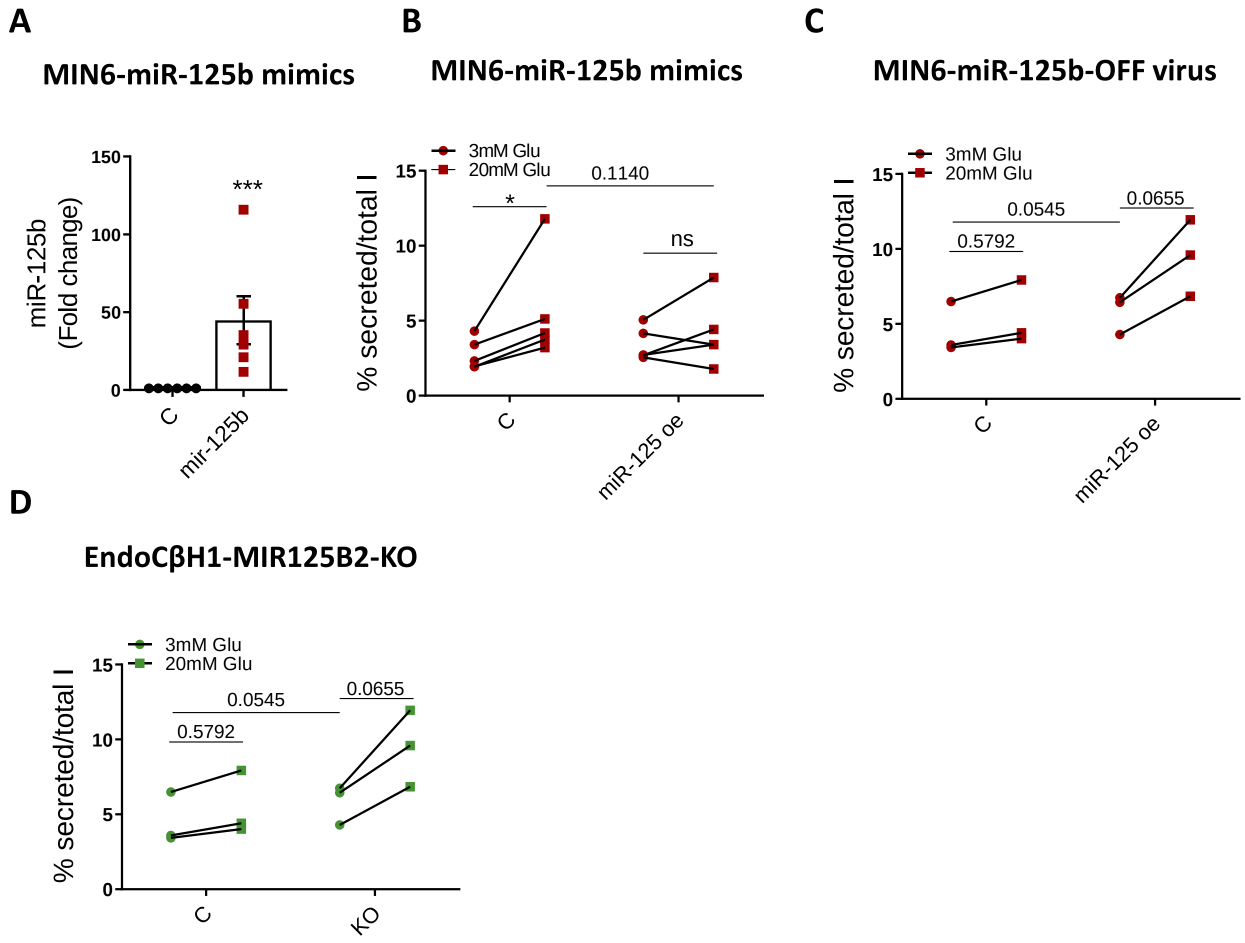

**Supplemental Figure 3**

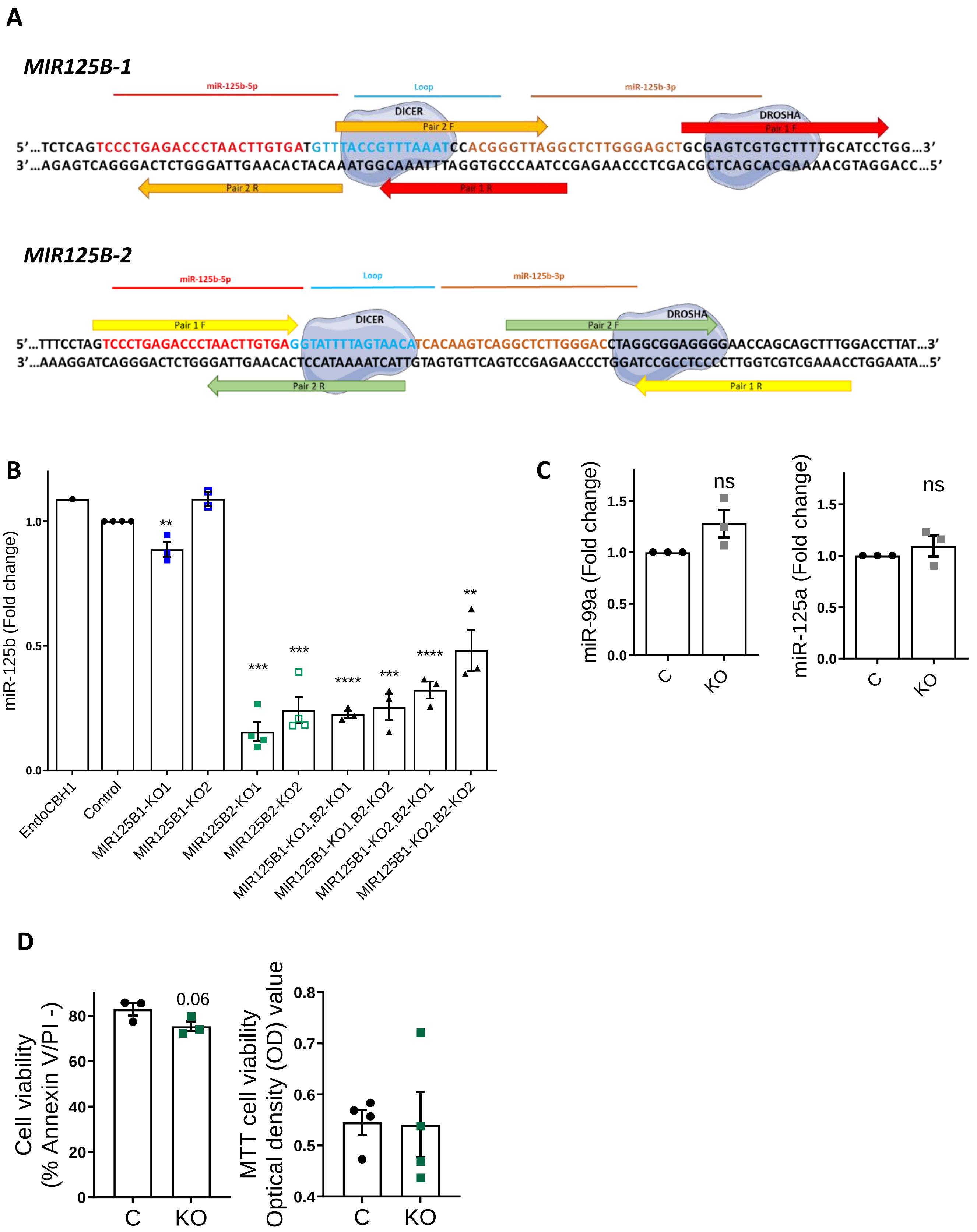

**Supplemental Figure 4.**

**A**

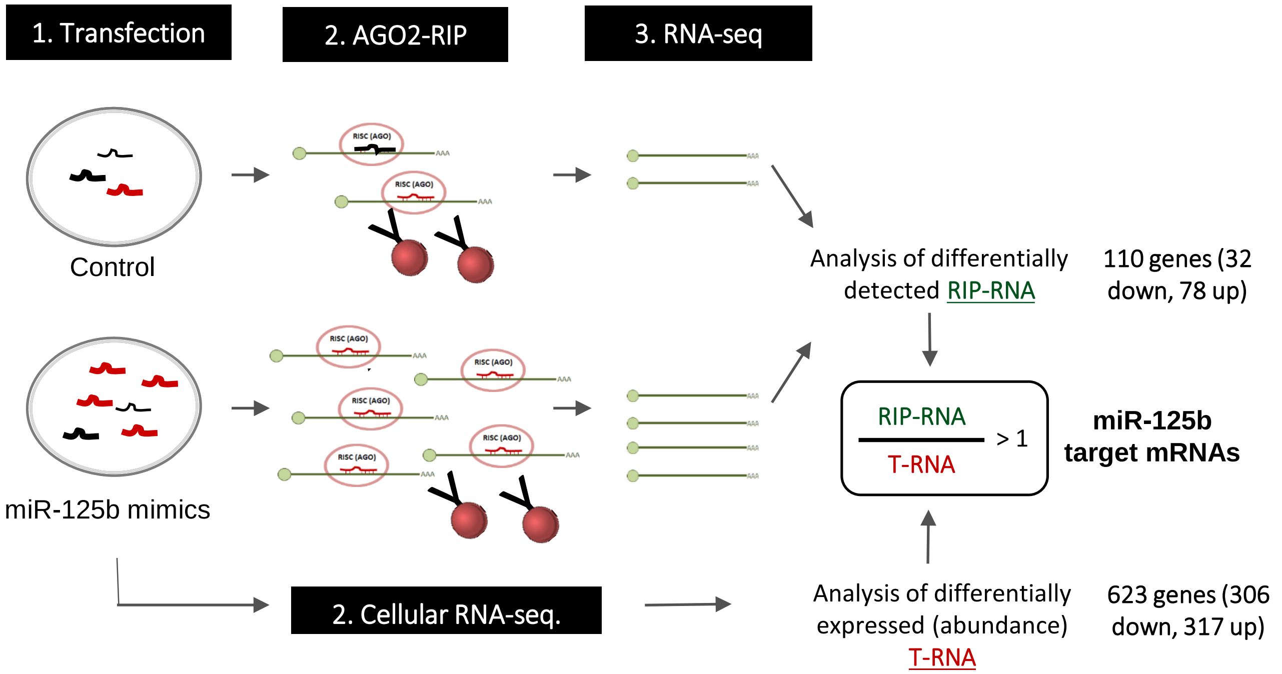


**B**

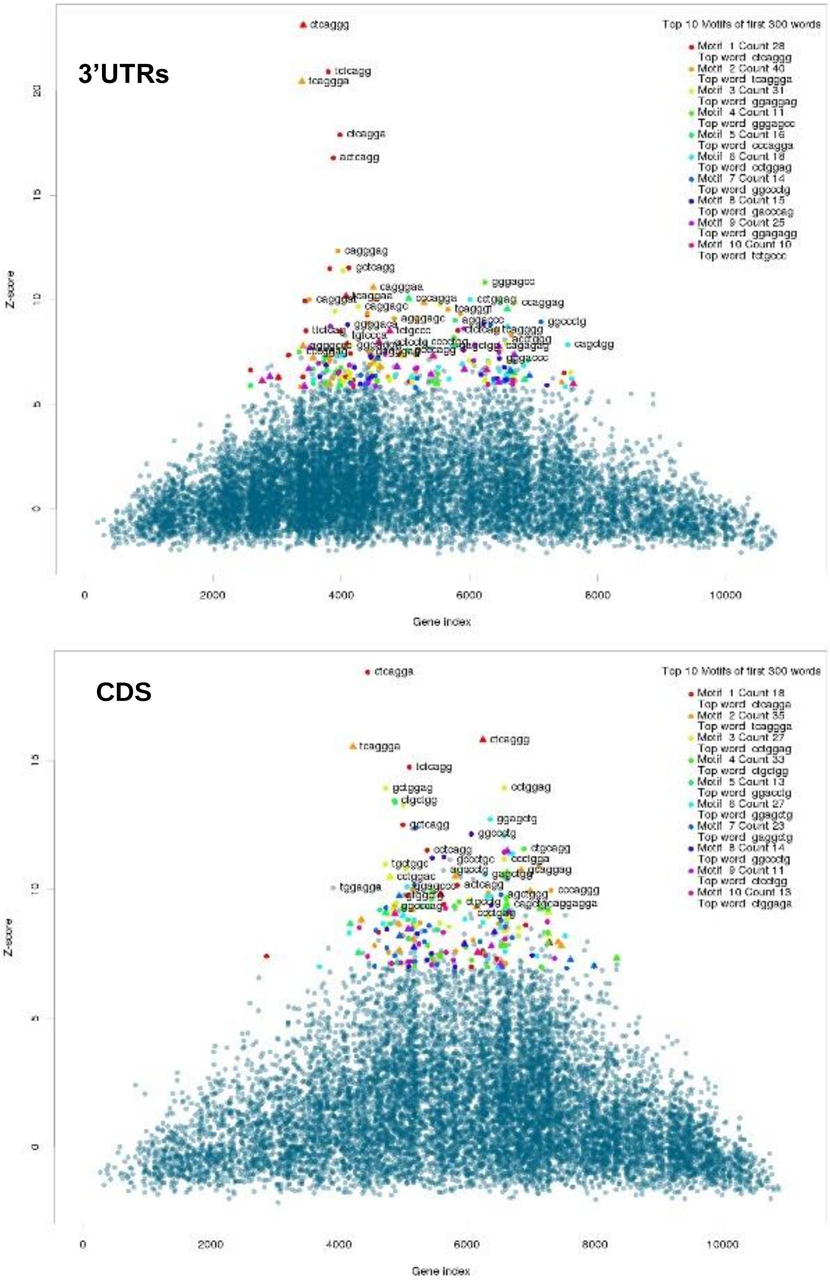


>mmu-miR-125b-5p
UCCCUGAGACCCUAA
CUUGUGA

RC: TCTCAGGGA

**Supplemental Figure 5**

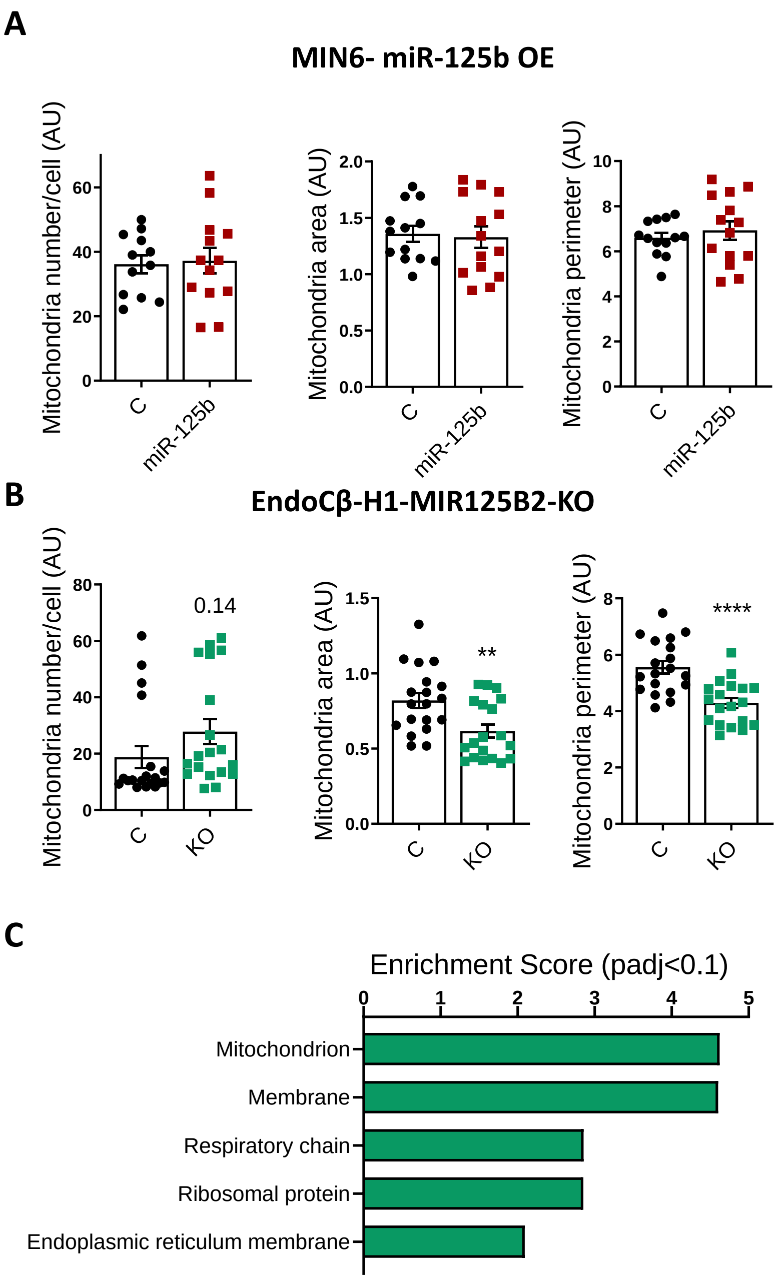

Supplemental Figure 6

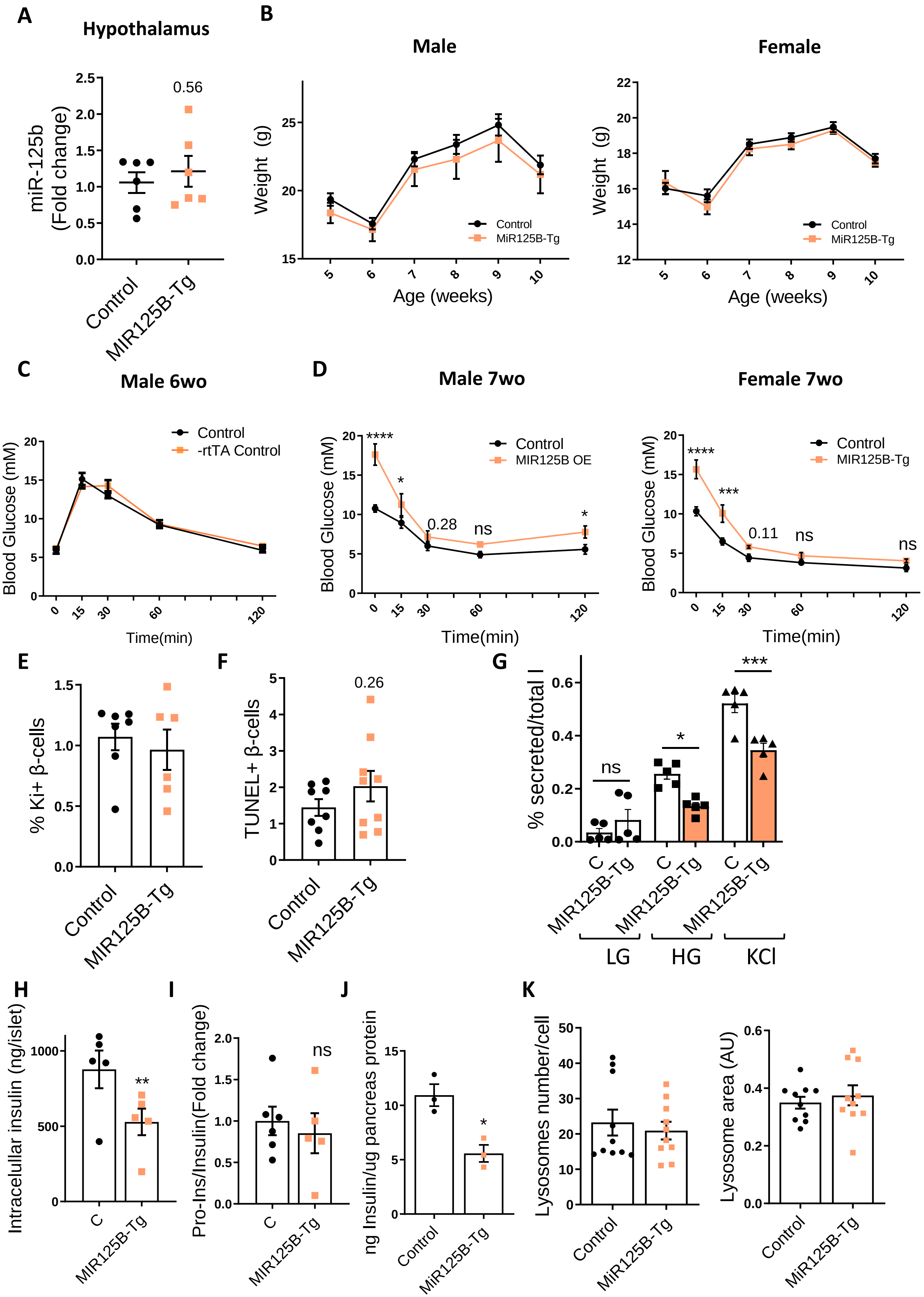

### Supplemental Figure 7

### A

Insulin secretion

| Gene | log2FoldChange |
|---|---|
| FFAR1 | -1.21974 |
| UCN3 | -1.19986 |
| GPR119 | -1.11149 |
| GIPR | -0.98636 |
| SLC2A2 | -0.94807 |
| SSTR3 | -0.9436 |
| MLXIPL | -0.86601 |
| GLP1R | -0.80968 |
| GLUL | -0.71175 |
| PDX1 | -0.67944 |
| KCNU1 | -0.6569 |
| PRKCB | -0.5862 |
| NKX6-1 | -0.58541 |
| CREB3L1 | -0.57256 |
| NEUROD1 | -0.47897 |
| SLC37A4 | -0.4743 |
| NKX2-2 | -0.43273 |
| AACS | -0.33216 |

Golgi

| Gene | log2FoldChange |
|---|---|
| EMID1 | -1.6699 |
| MSLN | -1.38883 |
| B3GALT5 | -1.07033 |
| B3GALNT1 | -0.9301 |
| CNTNAP2 | -0.88555 |
| GALNT14 | -0.84766 |
| MCFD2 | -0.83908 |
| ZDHHC2 | -0.82524 |
| PLK3 | -0.82351 |
| BMP1 | -0.77066 |
| GALNT18 | -0.75762 |
| PCSK9 | -0.69667 |
| CHPT1 | -0.68141 |
| EGFR | -0.6743 |
| SVIP | -0.65393 |
| PGAP3 | -0.57013 |
| RNF128 | -0.55974 |
| HEPACAM2 | -0.54779 |
| B3GNT3 | -0.54375 |
| DUSP26 | -0.53542 |
| PKDCC | -0.53344 |
| FUT10 | -0.51535 |
| GGA2 | -0.50119 |
| POFUT2 | -0.47267 |
| NMT2 | -0.46052 |
| LDLR | -0.45331 |
| TMEM167 | -0.44032 |
| KDELR1 | -0.43078 |
| TMED9 | -0.39473 |
| LMAN2 | -0.37793 |

Glycoproteins

| Gene | log2FoldChange |
|---|---|
| TMIGD3 | -2.00788 |
| CAR15 | -1.86875 |
| EMID1 | -1.6699 |
| TMIGD3 | -1.63581 |
| CDH7 | -1.52648 |
| FFAR3 | -1.40117 |
| MSLN | -1.38883 |
| VWA5B1 | -1.38188 |
| DCT | -1.37076 |
| TMED6 | -1.35608 |
| MUC4 | -1.25905 |
| CDHR1 | -1.22122 |
| SLC40A1 | -1.22104 |
| FFAR1 | -1.21974 |
| TMPRSS4 | -1.21359 |
| GCGR | -1.20314 |
| CPB2 | -1.10412 |
| B3GALT5 | -1.07033 |
| PRRT3 | -1.00494 |
| GIPR | -0.98636 |
| SLC2A2 | -0.94807 |
| SSTR3 | -0.9436 |
| B3GALNT1 | -0.9301 |
| CPM | -0.90018 |
| CNTNAP2 | -0.88555 |
| GPA33 | -0.88485 |
| CEACAM1 | -0.86556 |
| HCN4 | -0.85993 |
| ASIC1 | -0.84837 |
| SLC22A23 | -0.83433 |
| GLP1R | -0.80968 |
| ADORA1 | -0.80935 |
| OTOA | -0.80259 |
| PCSK4 | -0.79671 |
| IGFALS | -0.77858 |
| BMP1 | -0.77066 |
| PLA2G2F | -0.76601 |
| SYT2 | -0.75823 |
| FRAS1 | -0.75803 |
| GALNT18 | -0.75762 |
| PCSK9 | -0.69667 |
| GGT7 | -0.67943 |
| SLC18A1 | -0.67778 |
| EGFR | -0.6743 |
| OLFM4 | -0.63233 |
| FRZB | -0.62688 |
| TRAM1 | -0.62632 |
| G6PC2 | -0.61877 |
| PON2 | -0.61206 |
| NEU1 | -0.60585 |
| GUCY2C | -0.58323 |
| FAM174B | -0.5764 |
| CREB3L1 | -0.57256 |
| SLC7A5 | -0.5722 |
| PGAP3 | -0.57013 |
| RNF128 | -0.55974 |
| AMIGO2 | -0.55227 |
| HEPACAM2 | -0.54779 |
| B3GNT3 | -0.54375 |
| GALR1 | -0.5343 |
| PKDCC | -0.53344 |
| PIGO | -0.5264 |
| GRIN2C | -0.52029 |
| FUT10 | -0.51535 |
| TSPAN5 | -0.51337 |
| ADRA2A | -0.50873 |
| QSOX2 | -0.50759 |
| SIRPA | -0.4979 |
| TMEM231 | -0.4879 |
| POFUT2 | -0.47267 |
| LDLR | -0.45331 |
| NAGA | -0.44593 |
| GALNS | -0.44077 |
| TMEM167 | -0.44032 |
| PXMP4 | -0.43529 |
| TTYH2 | -0.43377 |
| TMEM8 | -0.42853 |
| LRFN4 | -0.42305 |
| SLC29A1 | -0.4132 |
| TMED9 | -0.39473 |
| INA | -0.38035 |
| LMAN2 | -0.37793 |
| GNS | -0.31404 |

ER

| Gene | log2FoldChange |
|---|---|
| EMID1 | -1.6699 |
| TMED6 | -1.35608 |
| MUC4 | -1.25905 |
| CLN6 | -1.10723 |
| B3GALT5 | -1.07033 |
| FMO1 | -1.01853 |
| SLC35D3 | -0.93592 |
| DIO1 | -0.90529 |
| MCFD2 | -0.83908 |
| ZDHHC2 | -0.82524 |
| ADORA1 | -0.80935 |
| ELOVL2 | -0.79347 |
| GRIP1 | -0.77811 |
| ESYT1 | -0.75418 |
| PCSK9 | -0.69667 |
| EGFR | -0.6743 |
| SVIP | -0.65393 |
| TRAM1 | -0.62632 |
| G6PC2 | -0.61877 |
| GUCY2C | -0.58323 |
| CREB3L1 | -0.57256 |
| PGAP3 | -0.57013 |
| RNF128 | -0.55974 |
| SEC31A | -0.54945 |
| PIGO | -0.5264 |
| LSS | -0.50743 |
| ANGEL1 | -0.50638 |
| POR | -0.49357 |
| ATP2A3 | -0.47299 |
| POFUT2 | -0.47267 |
| ATL2 | -0.44289 |
| JPH3 | -0.44126 |
| KDELR1 | -0.43078 |
| HS1BP3 | -0.40551 |
| TMED9 | -0.39473 |
| SEC61A1 | -0.39407 |
| INSIG1 | -0.35492 |

**Supplemental Figure 7**

**B**

Phosphoproteins

| Gene | log2FoldChange |
|---|---|
| PGM5 | 3.384586 |
| SRCIN1 | 2.143274 |
| CRMP1 | 1.994343 |
| SLC5A7 | 1.915163 |
| CDH6 | 1.903529 |
| STUM | 1.866269 |
| LSAMP | 1.856173 |
| NEFL | 1.735895 |
| NOS1 | 1.714182 |
| ELAVL2 | 1.708331 |
| ILDR2 | 1.683182 |
| FGFR2 | 1.626991 |
| CHL1 | 1.625863 |
| DOC2B | 1.57419 |
| ARHGAP6 | 1.569496 |
| DPYSL5 | 1.556181 |
| DPYSL3 | 1.537004 |
| MAPK4 | 1.528335 |
| MAP1A | 1.510319 |
| PCP4L1 | 1.467693 |
| FGFR3 | 1.453484 |
| DCX | 1.442323 |
| SH3PXD2B | 1.433606 |
| TIAM1 | 1.431407 |
| ADGRG6 | 1.431193 |
| PLCD3 | 1.425484 |
| EPN3 | 1.423582 |
| LEPR | 1.415607 |
| COL18A1 | 1.388956 |
| SYT1 | 1.382168 |
| KLHDC7A | 1.380444 |
| ADCYAP1R1 | 1.373177 |
| CARD10 | 1.365265 |
| FERMT1 | 1.35601 |
| SULT1A1 | 1.332353 |
| SOX5 | 1.322169 |
| DTNA | 1.313236 |
| ERBB2 | 1.297039 |
| CLDN2 | 1.285768 |
| DPP10 | 1.276844 |
| ATP1A3 | 1.274655 |
| ANO1 | 1.269384 |
| SFN | 1.249896 |
| REEP1 | 1.237442 |
| HSPB8 | 1.18827 |
| EVPL | 1.18489 |
| SHROOM1 | 1.183754 |
| KIF5A | 1.178635 |
| CLMN | 1.166565 |
| PYGB | 1.134098 |
| TUBB3 | 1.132731 |
| SDC1 | 1.128319 |
| BACH2 | 1.128003 |
| KIRREL | 1.122798 |
| AMOT | 1.110241 |
| DCLK1 | 1.110144 |
| KCNA6 | 1.107624 |
| STRIP2 | 1.102385 |
| KCNN4 | 1.099209 |
| SLC52A3 | 1.098478 |
| HK1 | 1.093945 |
| WWC1 | 1.085285 |
| ARNTL | 1.083724 |
| PLCE1 | 1.081234 |
| PALMD | 1.078257 |
| KCNQ1 | 1.075632 |
| PTPN14 | 1.064004 |
| MMP14 | 1.0492 |
| CNR1 | 1.044278 |
| HMGCS2 | 1.043029 |
| KCNQ2 | 1.039918 |
| GAS6 | 1.032908 |
| PLS3 | 1.031522 |

Axon

| Gene | log2FoldChange |
|---|---|
| SRCIN1 | 2.143274 |
| SLC5A7 | 1.915163 |
| NEFL | 1.735895 |
| NTM | 1.73145 |
| CALB1 | 1.523657 |
| SYT1 | 1.382168 |
| DTNA | 1.313236 |
| HTR3A | 1.286413 |
| ATP1A3 | 1.274655 |
| KIF5A | 1.178635 |
| NCS1 | 1.150788 |
| PYGB | 1.134098 |
| TUBB3 | 1.132731 |
| KCNA6 | 1.107624 |
| TGFB2 | 1.089331 |
| CNR1 | 1.044278 |
| SNCA | 1.021912 |
| PALLD | 1.01456 |
| ACHE | 0.974595 |
| KCNA2 | 0.881891 |
| SEMA6A | 0.765375 |
| PALM | 0.764501 |
| ALCAM | 0.751319 |
| MAPT | 0.723769 |
| KCNH1 | 0.683248 |
| MYH10 | 0.679126 |
| CTNNA2 | 0.629709 |
| APP | 0.491763 |
| CPT1C | 0.468435 |
| MAP2K1 | 0.348694 |

Glycoproteins

| Gene | log2FoldChange |
|---|---|
| KCNK13 | 3.078916 |
| SMOC1 | 3.040074 |
| GABRP | 2.117097 |
| PDGFC | 2.040732 |
| WNT7B | 2.036377 |
| DLK1 | 1.917689 |
| SLC5A7 | 1.915163 |
| CDH6 | 1.903529 |
| EVC2 | 1.875716 |
| AQP5 | 1.864548 |
| LSAMP | 1.856173 |
| COL27A1 | 1.820278 |
| NEFL | 1.735895 |
| NTM | 1.73145 |
| P3H2 | 1.6516 |
| FGFR2 | 1.626991 |
| CHL1 | 1.625863 |
| FLRT3 | 1.616883 |
| TNFRSF19 | 1.564367 |
| TGFB3 | 1.547986 |
| MFAP2 | 1.547205 |
| KL | 1.482683 |
| SCNN1A | 1.469131 |
| SCARA3 | 1.465067 |
| FGFR3 | 1.453484 |
| ADGRG6 | 1.431193 |
| ST6GALNAC2 | 1.424805 |
| LEPR | 1.415607 |
| FLRT2 | 1.413927 |
| COL18A1 | 1.388956 |
| SYT1 | 1.382168 |
| KLHDC7A | 1.380444 |
| ADCYAP1R1 | 1.373177 |
| COL9A2 | 1.358039 |
| INHBA | 1.342248 |
| OLFML2B | 1.305021 |
| ERBB2 | 1.297039 |
| HTR3A | 1.286413 |
| SPOCK3 | 1.280884 |
| DPP10 | 1.276844 |
| ANO1 | 1.269384 |
| CD14 | 1.25192 |
| CP | 1.229932 |
| PRELP | 1.209164 |
| LAMA5 | 1.189775 |
| SNED1 | 1.153812 |
| LYNX1 | 1.1496 |
| GDF10 | 1.140987 |
| SDC1 | 1.128319 |
| APCDD1 | 1.128 |
| KIRREL | 1.122798 |
| ST3GAL4 | 1.101342 |
| LGI2 | 1.099607 |
| SLC52A3 | 1.098478 |
| TGFB2 | 1.089331 |
| ARNTL | 1.083724 |
| ADAMTSL4 | 1.083002 |
| KCNQ1 | 1.075632 |
| SEMA3B | 1.073551 |
| TMEM59L | 1.052226 |
| SLIT2 | 1.051137 |
| RNF43 | 1.046204 |
| CNR1 | 1.044278 |
| GAS6 | 1.032908 |
| PLAUR | 1.030368 |
| PLAT | 1.02416 |
| AGRN | 1.020408 |
| NOTCH2 | 1.013127 |
| ENPP3 | 1.011407 |
| HAPLN1 | 1.011075 |
| SEMA3C | 1.009993 |
| SEMA4F | 1.007728 |
| CRIM1 | 1.003023 |

Golgi

| Gene | log2FoldChange |
|---|---|
| P3H2 | 1.6516 |
| FGFR2 | 1.626991 |
| SCARA3 | 1.465067 |
| FGFR3 | 1.453484 |
| EHF | 1.447345 |
| ST6GALNAC2 | 1.424805 |
| MAL | 1.395117 |
| SYT1 | 1.382168 |
| ATP1A3 | 1.274655 |
| CD14 | 1.25192 |
| HSPB8 | 1.18827 |
| NCS1 | 1.150788 |
| ST3GAL4 | 1.101342 |
| PLCE1 | 1.081234 |
| TMEM59L | 1.052226 |
| RAB27B | 1.049924 |
| SNCA | 1.021912 |
| ACHE | 0.974595 |
| APC2 | 0.974186 |
| ST6GALNAC2 | 0.973201 |
| IL17RD | 0.969734 |
| PTGS1 | 0.958713 |
| SMO | 0.944786 |
| ECE2 | 0.943918 |
| PLD1 | 0.928606 |
| CDC42EP1 | 0.843105 |
| PTGFRN | 0.841552 |
| CAV2 | 0.825733 |
| PARM1 | 0.821136 |
| MDFIC | 0.808626 |
| GJA1 | 0.773144 |
| EMP2 | 0.77238 |
| SMPD3 | 0.70152 |
| CREG2 | 0.699238 |
| SORL1 | 0.657576 |
| IRGM2 | 0.639081 |
| LARGE1 | 0.63514 |
| SPRY1 | 0.627566 |
| ZDHHC9 | 0.614026 |
| RAB31 | 0.612486 |
| APP | 0.491763 |
| ENTPD6 | 0.398712 |
| MAP2K1 | 0.348694 |

### Supplemental Figure 8

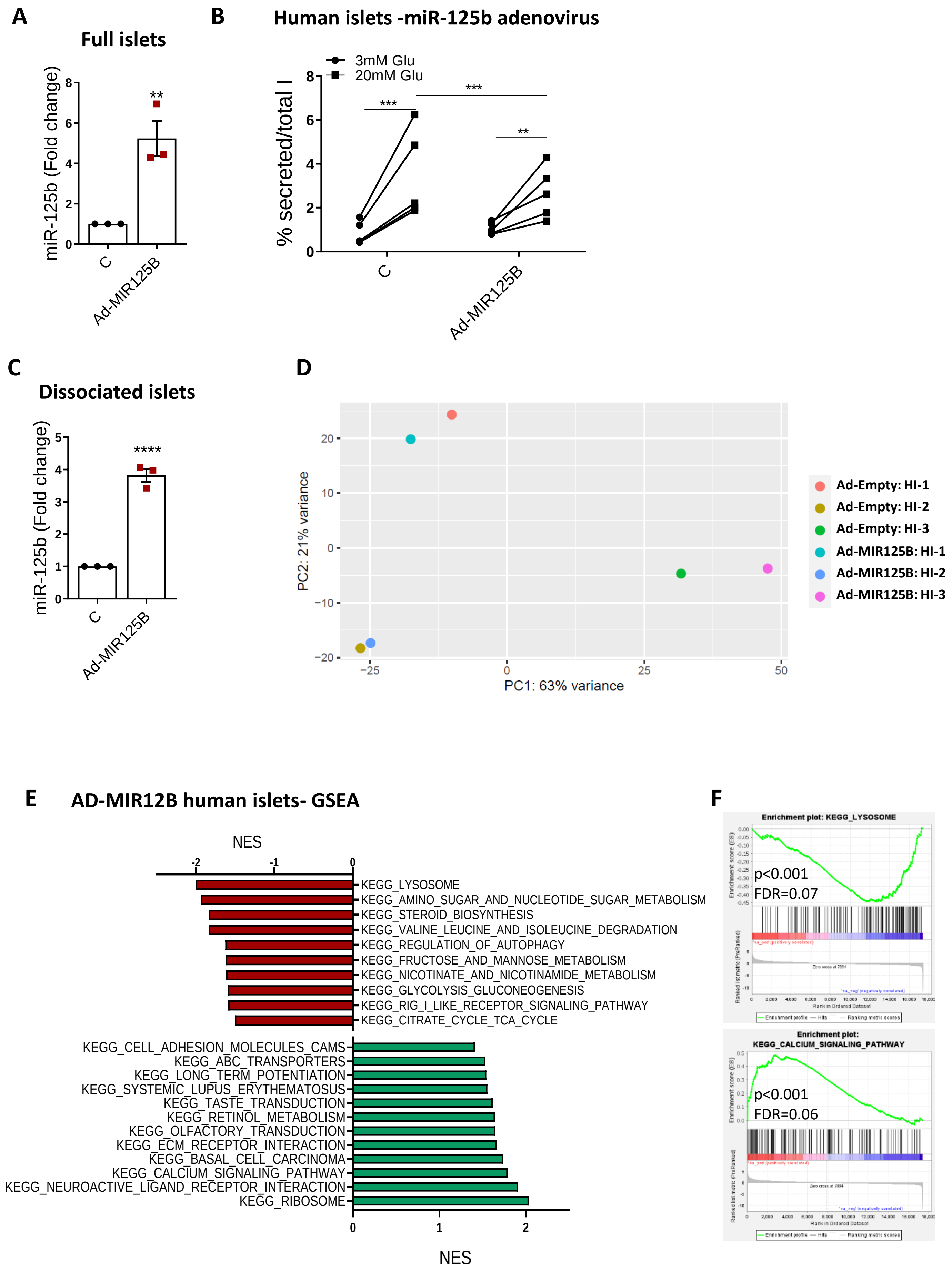
